# The Phage Time Continuum: Genome Organization Constrains Anti-CRISPR Success

**DOI:** 10.64898/2026.10.15.759504

**Authors:** Sean Czerwinski, Iris Irby, Gabriel Morales Artavia, Sam P Brown, James Gurney

## Abstract

Bacteria and lytic phages experience antagonistic co-evolution, with bacterial defenses (e.g., CRISPR-Cas) battling phage anti-defenses (e.g., anti-CRISPR (Acr) systems). We investigate whether CRISPR-Cas and Acr engage in a timing-dependent genetic conflict. If anti-CRISPR genes act too late in infection, they risk failing to neutralize CRISPR, if Cas cannot find its target in time, Acr may be able to neutralize it. Because phages typically inject their genomes in a linear manner, location along the genome acts as a proxy for time from start of infection. Given these restrictions, we suggest that phage genomes that contain Acr systems will have a distinct pattern of spacers when compared to those that do not, reflecting the time dependent interaction of infections, we have termed this the Phage Time Continuum. In our analysis of 1462 phage genomes and 105 mobile plasmid genomes infecting *Pseudomonas aeruginosa*, we found clear distribution of where *P. aeruginosa* spacers map back onto these mobile genetic elements’ genomes. Further CRISPR spacers significantly enriched against anti-CRISPR systems over other anti-defense systems from both phage and plasmids. Finally, in genomes with anti-CRISPR, spacers map to earlier positions, suggesting that success hinges not just on targeting a given gene, but where those spacers hit along a genome.

## Introduction

CRISPR-Cas systems are bacterial adaptive immune systems that target foreign nucleic acids [1]. These systems act as genetic immune memories, incorporating short stretches of invading genetic material from mobile genetic elements (MGEs, e.g., phages, plasmids and transposons) into specific host loci called CRISPR arrays [2-4]. In doing so, hosts form a genetic memory of past infections [5, 6]. Spacers are transcribed into CRISPR RNAs (crRNA) which guide CRISPR-associated (Cas) proteins to scan intracellular nucleic acids for matching sequences and in the event of sequence alignment, cleave the target nucleic acid [2, 7, 8]. Much is known about molecular mechanisms of Cas proteins and their use as genetic tools for DNA cleavage; however, the CRISPR spacer acquisition step (adaptation) has proven more difficult to understand [9-11]. CRISPR adaptation has been rarely observed in real-time, and the factors that constrain spacer acquisition across an invading genome remain incompletely understood [9, 12-14]. Considering that many tailed dsDNA phages inject their genome linearly, including cos- and pac/headful packaging phages, genomic position relative to the first gene to enter may therefore act as a proxy for time along infection, producing a time continuum [15, 16]. Similarly, during conjugation, mobilizable plasmids are transferred directionally as linear ssDNA, so location relative to *oriT* is a proxy for time since entry [17-21]. While filamentous phages enter as circular ssDNA the relevant temporal constraint may arise from transcription-dependent DNA topology rather than directional genome entry [22, 23]. Negative supercoiling generated behind elongating RNA polymerase can facilitate Cas R-loop formation and subsequent CRISPR target cleavage[24-27]. Thus, genomic regions transcribed earlier may be preferentially accessible to CRISPR-Cas surveillance. This physical property of many MGEs may apply an evolutionary pressure on gene synteny.

In addition to temporal structure, some MGEs carry anti-CRISPR (Acr) systems that inhibit the function of Cas proteins [28]. These antidefense systems have been shown to be commonly encoded in the leading regions of mobilizable plasmid genomes [18]. In phages, Acr genes are often expressed early during infection; however, their genomic position relative to the first-entering end of the injected genome has not, to our knowledge, been systematically evaluated [29, 30]. The positioning observed in plasmids makes sense if the efficacy of early anti-defense gene products are dependent on their rapid deployment. In the case of Acr, early expression will increase the likelihood of inhibiting Cas activity [28, 29]. If Acr genes act too late in infection, they risk failing to neutralize Cas before it can find PAMs and degrade the MGE’s nucleic acid [31, 32]. Thus, these two systems may engage in a timing-dependent genetic conflict which we have termed the Phage Time Continumm (Figure 1).

**Figure 1.**
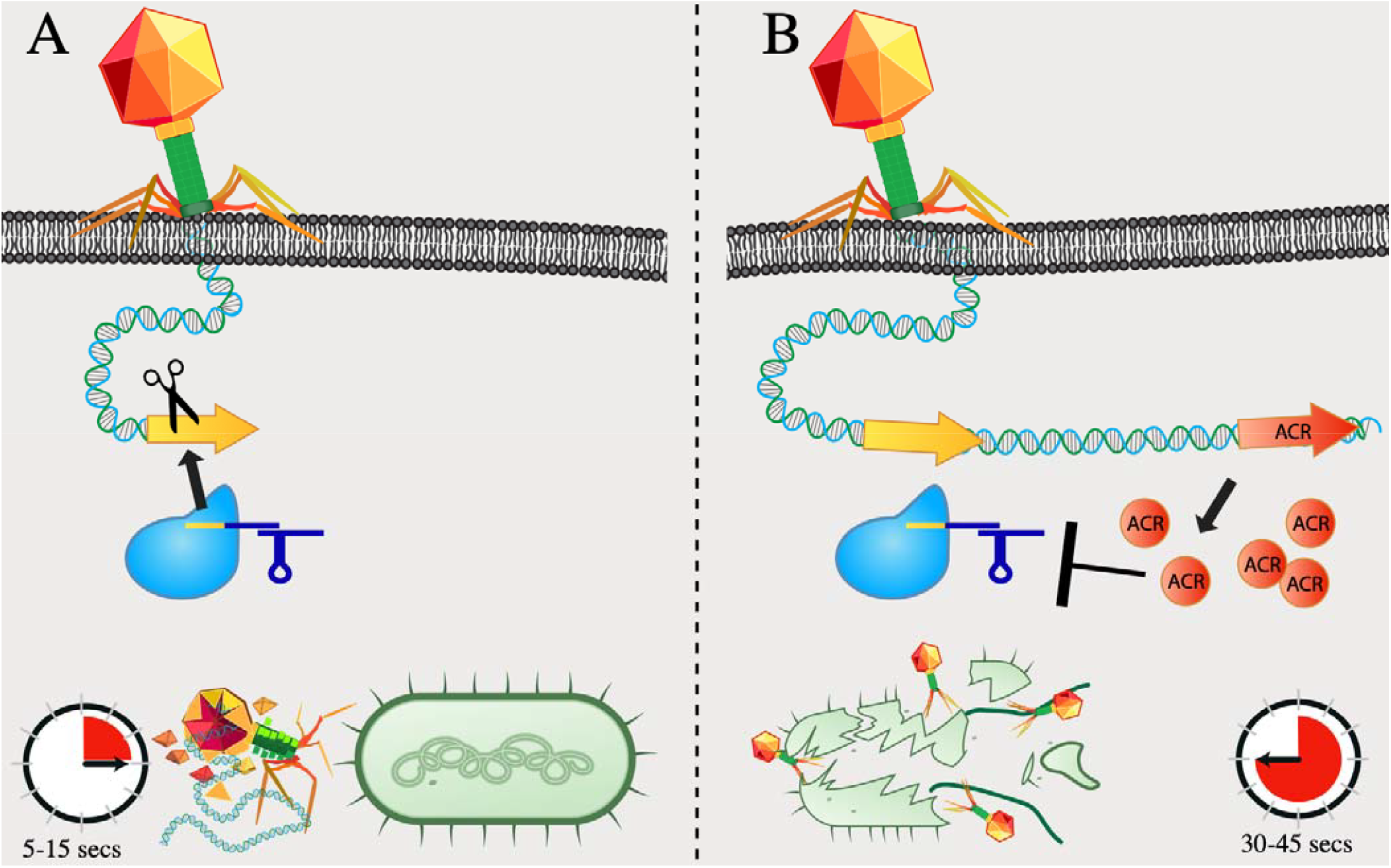
Schematic illustration of the Phage Time Continuum hypothesis. Phage genomes that enter in a linear manner have a time dimension in regard to when CRISPR “sees” each gene. CRISPR-Cas is shown in blue, the CRISPR target/spacer in yellow, and Acr in red. A) The CIRSPR system is targeting an early gene in the phage genome. This leads to the bacteria surviving. B) the CRISPR system is targeting a phage gene that is delivered after an Anti-CRISPR gene, leading to the inhibition of CRISPR and the phage lysing the cell.

The temporal nature of MGE infection suggests three alternate hypotheses for the distribution of CRISPR spacers along invading genomes. Under our exposure-only null hypotheses, spacer representation primarily reflects how often a target sequence is encountered, with no additional effect of genome position. In contrast, our general phage time continuum hypothesis states that target sequences entering earlier should be preferentially represented because they are available to to CRISPR sooner, regardless of Acr status. Finally, our Acr-dependent phage time continuum hypothesis posits that the early-targeting bias is strengthened specifically in Acr+ phages, because Acr shortens the window for effective CRISPR targeting. These hypotheses therefore predict, respectively, no positional bias, a general early-targeting bias, or an Acr-dependent enhancement of that bias.

To reveal the biases of CRISPR targeting, we must investigate the functions, locations, and genomic context of their targets. Using spacer sequences from *P. aeruginosa*, we mapped CRISPR targets on to 1462 phage and 105 plasmid genomes and examined their relationship to anti-defense genes. Our analysis reveals several striking results. First, there is a significant signature of CRISPR spacers being enriched for Acr systems relative to other MGE anti-defense systems. Second, these protospacer targets also appear significantly earlier in Acr-encoding parasites than in those without Acr systems, providing support for a time-based constraint when an infecting genome can counter CRISPR. Finally, spacers were found to be more likely than expected to hit anti-defense genes outside of the actual anti-defense distribution, suggestive of a broader implication for the role gene synteny may play in response to defense systems. These results suggest that CRISPR-Cas targeting is constrained by both targeting genes with specific function as well as overall genomic architecture in the form of where those genes fall within the genome of the MGE.

## Results and Discussion

### Anti-CRISPR systems are disproportionately targeted by CRISPR spacers

The spacer record is an inherently incomplete archive of encounters with MGEs. First, not all [12, 33, 34] infections produce spacers. Although a spacer may be acquired during a given infection, it will only persist in the record if the infected cell and its progeny survive[35, 36]. Thus, lytic infections that proceed to host lysis or chronic prophages that targeting would result in autoimmunity, may be underrepresented in spacer records, producing a survivorship bias toward spacers associated with successful immunity[15, 34, 37, 38]. Second, there must be continued selection for a given spacer to remain relevant, as genetic drift and selection for other spacers erase or rewrite its presence making the array a biological palimpsest[39]. As a result, some encounters are inscribed, some subsequently erased, and many leave no trace. If Acr are highly effective at inhibiting CRISPR then spacers which target Acrs could be rare. However, if there is sufficient time between genome injection, CRISPR-cas PAM searching and the Acr system’s attempt to deactivate CRISPR would could see disproportionately high spacer hit rates compared to other anti-defense classes. If the Phage Time Continuum model holds – Acr should be enriched early in phage genomes, relative to injection, and CRISPR spacers should preferentially target early phage genes regardless of function.

Constraining our results to spacers targeting anti-defense systems (a total of 7162 spacers), Acr systems are disproportionately targeted. In phages, an average Acr system received 42.7 spacer hits versus 1.8 for other anti-defense systems (∼24× enrichment; SI Table 1); in plasmids, 4.2 versus 0.5 (∼8×; SI Table 1). Across both MGEs, ∼94% of anti-defense-targeting spacer hits fell on Acr (6,740 Acr vs 422 other in phages; 851 Acr vs 55 other in plasmids), despite Acr representing 39.6% of phage anti-defense systems and 66.0% of plasmid anti-defense systems. Most other anti-defense classes are near zero or very low (Figure 2, which holds even when normalized for the number of those systems in each phage **SI Figure 1**). In phage, only anti-CBASS (anti-Cyclic oligonucleotide-Based Anti-phage Signaling System) and anti-RM (anti-Restriction Modification) were also targeted, and in plasmid only anti-RM. Using a generalized linear model where we controlled for the early enrichment of Acr genes we still find a significant increase in the abundance of spacers matching back to Acr genes. These data suggest that we can exclude our first hypothesis, the abundance of a given gene does not fully predict its relative abundance in the CRISPR-Cas spacer array.

**Figure 2.**
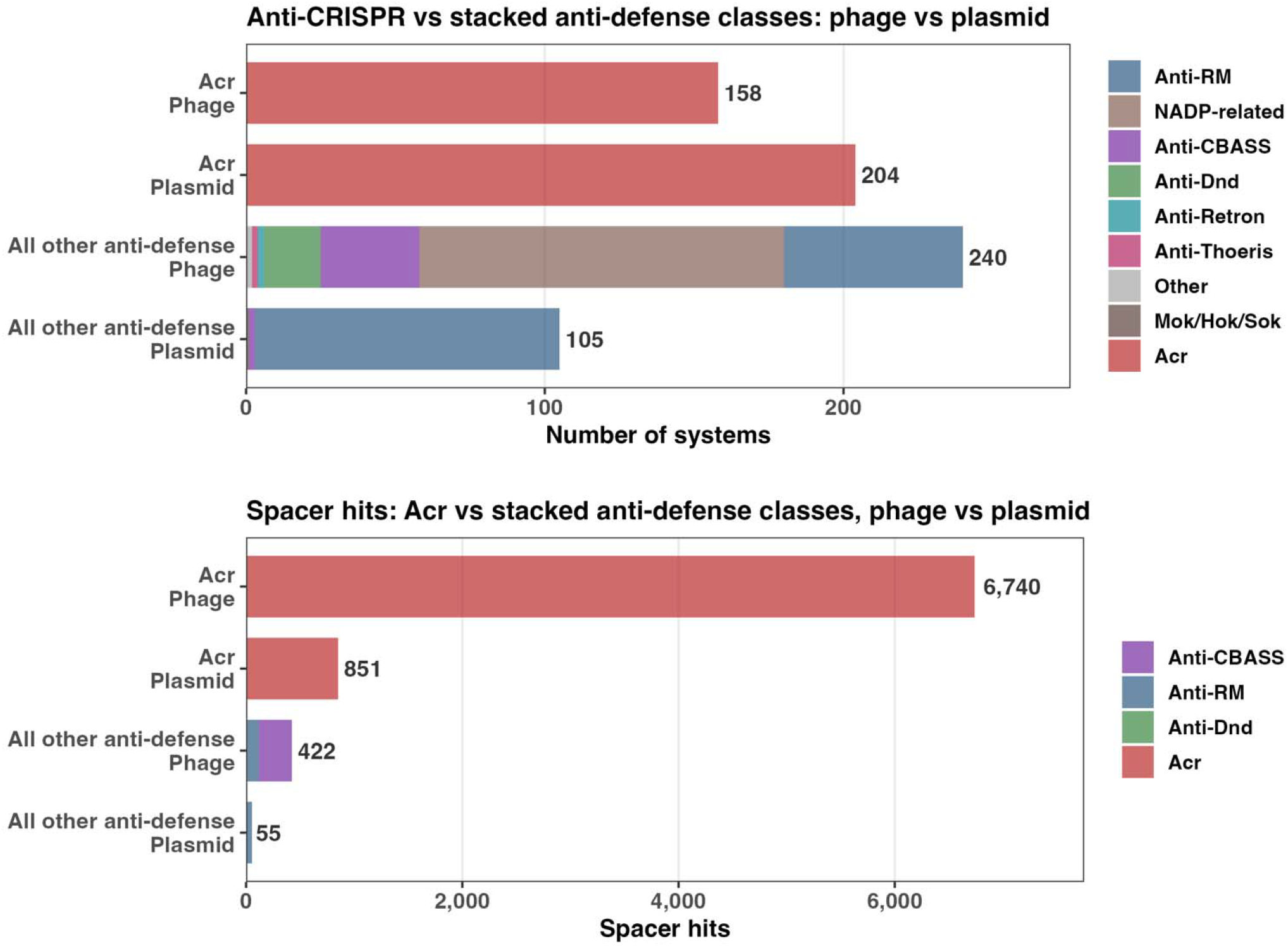
Anti-CRISPR (Acr) systems are disproportionately targeted by CRISPR spacers relative to other anti-defense systems. Anti-defense systems identified by DefenseFinder were grouped by class for phage and plasmid genomes, and CRISPR spacer hits overlapping each class were quantified. Bars show the number of anti-defense systems and the number of spacer hits associated with each class. Acr systems account for the large majority of anti-defense-targeting spacer hits in both phages and plasmids despite representing only a subset of the anti-defense systems present. In phages, Anti-CBASS and Anti-RM systems were also targeted, while Anti-RM was the only additional anti-defense class targeted in plasmids.

### Anti-defense systems and spacer targets occupy distinct genomic positions

There is additionally a clear pattern of CRISPR targeting phages with anti-CRISPR. All 98 Acr+ phage in our dataset were targeted by at least one CRISPR spacer, versus only 42.1% of Acr- phages (574 of 1364). The majority of phage (790/1462) had no spacers matches but all lacked Acr. Using genus and family level information, we do not see a clear picture of CRISPR targeting an isolated phage group (**SI Figure 2; SI table 2**) reducing our concern that the effect is taxonomical. The number of spacers that matched were also clearly disproportional; of all spacer blast hits, more than half (56.3%) mapped back to Acr+ phage. Given that Acr+ phage makes up only 98 of 1462 total genomes (6.7%), this is clearly significant (Binomial test p < 2x10^-16^). The presence of Acr+ substantially increases the likelihood of phages nucleic acid ending up in a spacer array.

To begin to interrogate our other two hypothesis we took a more detailed look at spacers mapping back to anti-defense systems. In phages, Acr genes are enriched within the first 20% of the normalized genome, and spacer hits largely follow the expected density (Figure 3A). This is contrasted by the mid-to-late Acr position in our plasmid dataset that departs from previous work, from which we derived our normalization method, that showed anti-defense genes generally enrich early in plasmids [18] (Figure 3B). Interestingly, in phages there is a significant signature of spacers mapping to regions outside of the expected density of the anti-RM and anti-CBASS systems (Figure 3C). In both cases this mapping occurred ahead of the expected density, once again suggesting an arms race towards earlier positioning. However, the distribution of spacers mapping back to anti-RM in plasmids is predicted by the distribution of those systems (Figure 3D). These data provide evidence for both hypotheses two and three, however we are yet to consider the presence or absence of anti-defense Acr systems.

**Figure 3.**
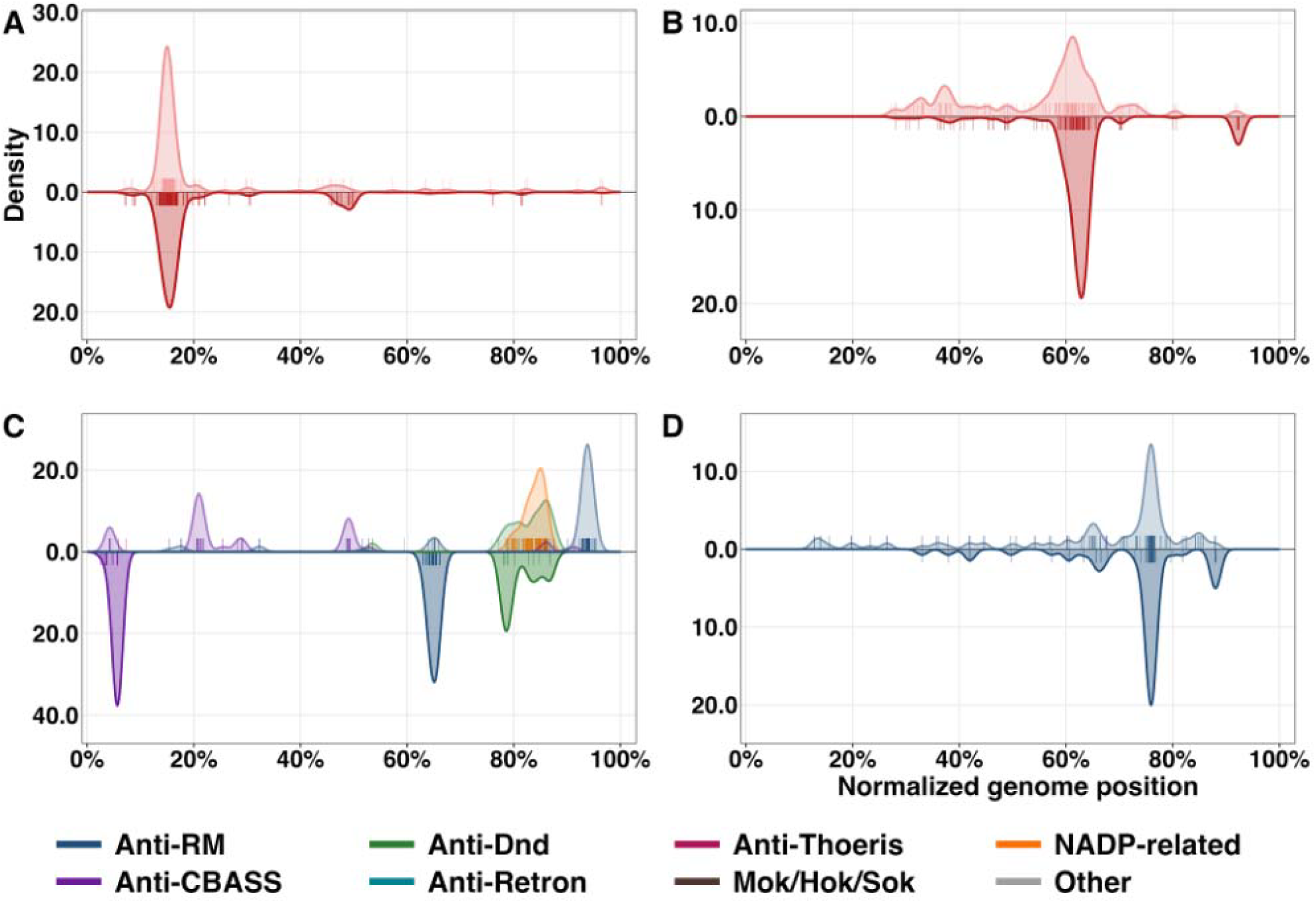
Genomic distributions of anti-defense systems and CRISPR spacer hits targeting those systems in phages and plasmids. Anti-defense systems were identified using DefenseFinder and their positions were mapped along normalized phage and plasmid genomes. For each panel, the upper distribution shows the normalized genomic positions of the indicated anti-defense systems, while the lower mirrored distribution shows the positions of CRISPR spacer hits overlapping those systems. A) Anti-CRISPR systems in phages. B) Anti-CRISPR systems in plasmids. C) non-Acr anti-defense systems in phages, including Anti-RM and Anti-CBASS systems targeted by spacers. D) non-Acr anti-defense systems in plasmids, where spacer targeting was restricted primarily to Anti-RM systems. Rug marks indicate the observed positions contributing to each distribution, and density curves are normalized within each category.

### Anti-CRISPR phages are targeted earlier in the genome

To try and disentangle hypotheses two and three we next asked a question of both phages and plasmids: does the genomic position of spacer hits differ between MGEs that carry Anti-CRISPR systems (Acr+) and those that do not (Acr-)? Spacers targeting Acr+ phages should be enriched toward earlier positions in the phage genome as Acrs are enriched early in the genomes. However, does this hold true for all the spacers that target these genomes? Our Phage Time Continuum hypothesis states earlier-targeting spacers would be selectively retained, as they can target positions available before Acr expression can inhibit Cas.

To test whether Acr presence was associated with a genome-wide shift in CRISPR targeting, rather than only with direct targeting of Acr loci, we compared the normalized positions of all retained spacer hits between Acr+ and Acr- phages. All 1,037,647 filtered spacer BLAST hits were pooled by Acr status to compare their normalized positions across Acr+ and Acr− phage genomes. Hits specifically to Acr genes were removed as we know that thse genes cluster near the start of phage genomes. Spacer hits on Acr+ phages occurred significantly earlier than those on Acr− phages (median position 44.9% vs 52.3%; one-sided Wilcoxon rank-sum test; Hodges-Lehmann shift = −0.049) (Figure 4; SI Figure 3; **SI Table 3**). Because this pooled comparison gives greater weight to phages with many spacer hits, we also performed a genome-level analysis in which each phage was represented by the median normalized position of all spacer hits mapping to that genome.

**Figure 4.**
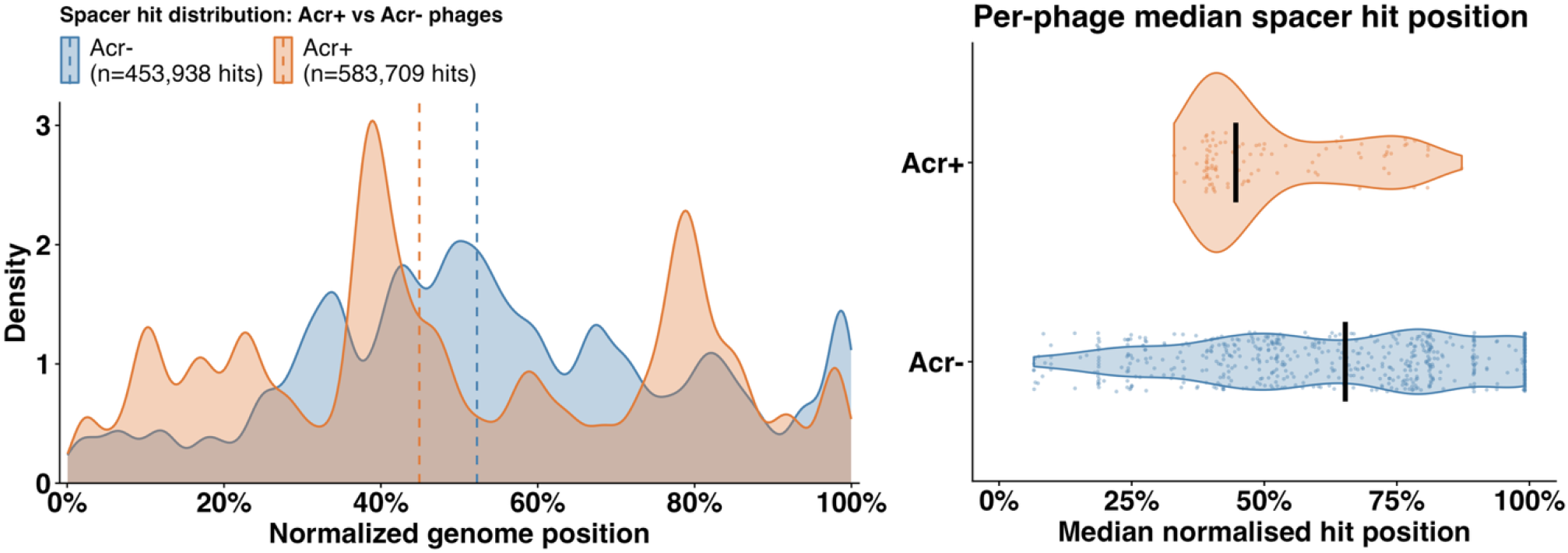
CRISPR spacer hits occur earlier in phage genomes carrying anti-CRISPR systems. Phage genomes were classified as Acr+ if they contained at least one Acr system identified by DefenseFinder and Acr- if no Acr system was detected. **Left)** Density distributions of spacer-hit positions along normalized phage genomes for Acr+ and Acr- phages. Dashed lines indicate the median spacer-hit position for each group. **Right)** Per-phage median spacer-hit positions. Spacer hits were first pooled within each phage, and the median normalized position was calculated for that genome so that each point represents one phage. Black crossbars indicate the median of the per-phage values within each group. Spacer targeting is shifted toward earlier genomic positions in Acr+ phages relative to Acr- phages.

On a per-phage basis, Acr+ phages were targeted earlier than Acr- phages, with median per-phage positions of 44.6% and 65.3%, respectively. The estimated positional shift was 12.8 percentage points toward the beginning of Acr+ genomes (Hodges-Lehmann shift = -0.128; one-sided Wilcoxon rank-sum test, p = 3.74 × 10□ □; **SI Table 4**). Because each targeted phage contributed a single observation, this comparison reduced the influence of heavily targeted genomes and provided a more conservative estimate of the positional shift. The pooled distribution showed the same directional pattern, with the first quartile of Acr+ spacer hits occurring by 28.1% of genome length compared with 37.3% for Acr- hits.

To determine whether this pattern was driven primarily by spacers directly targeting Acr loci, we repeated the analysis after removing spacer hits whose midpoint fell within annotated Anti-CRISPR system coordinates. Excluding these hits produced little change in the overall distribution or genome-level effect, indicating that the earlier targeting of Acr+ phages was not explained solely by the genomic positions of Acr loci themselves.

Together, these results support our third hypothesis that the retained spacer distribution reflects an interaction between genomic position and competition with anti-CRISPR systems. Spacer targets occur earlier in Acr+ phages, consistent with stronger selection for CRISPR targets that become available before Acr-mediated inhibition can be established.

### Anti-CRISPR plasmids show the same directional shift in spacer targeting

To generalize our data, we extended our analysis to plasmids as they serve as a parallel to phages. The life histories of phages and plasmids are different. We observed differences between phage and plasmids in terms of spacers and presence of anti-defense genes. First, a far larger proportion of plasmids carry an Acr gene than phages (72% in comparison to 6.7% in phage). As in phages, Acr+ plasmids are more frequently targeted than Acr-. All 76 Acr+ plasmids were targeted by at least one spacer versus 22 of 29 Acr- plasmids (75.9%; Fisher exact p=6.86x10^-5^). This result again supports the co-evolutionary interpretation: plasmids that are frequently targeted by CRISPR are the ones that evolve Acr countermeasures.

We next investigated if there was a similar distribution of spacers due to the presence or absence of Acr genes. All 176,914 (167,953 Acr+ / 8,961 Acr-) plasmid spacer analyses were computed identically to the phage analysis.

Again, groups were compared via kernel density estimation and Wilcoxon rank-sum test. The per-plasmid analysis shows that spacer targeting in Acr+ plasmids follows the same directional pattern observed in Acr+ phages. Interpreting genome position relative to the end that first enters the recipient cell, spacer hits on Acr+ plasmids are shifted toward the incoming end relative to Acr- (median 23.5% vs 35.5%; per-plasmid Hodges– Lehmann shift = −0.175, p = 1.95×10□ □; Figure 5; SI Table 5).

**Figure 5.**
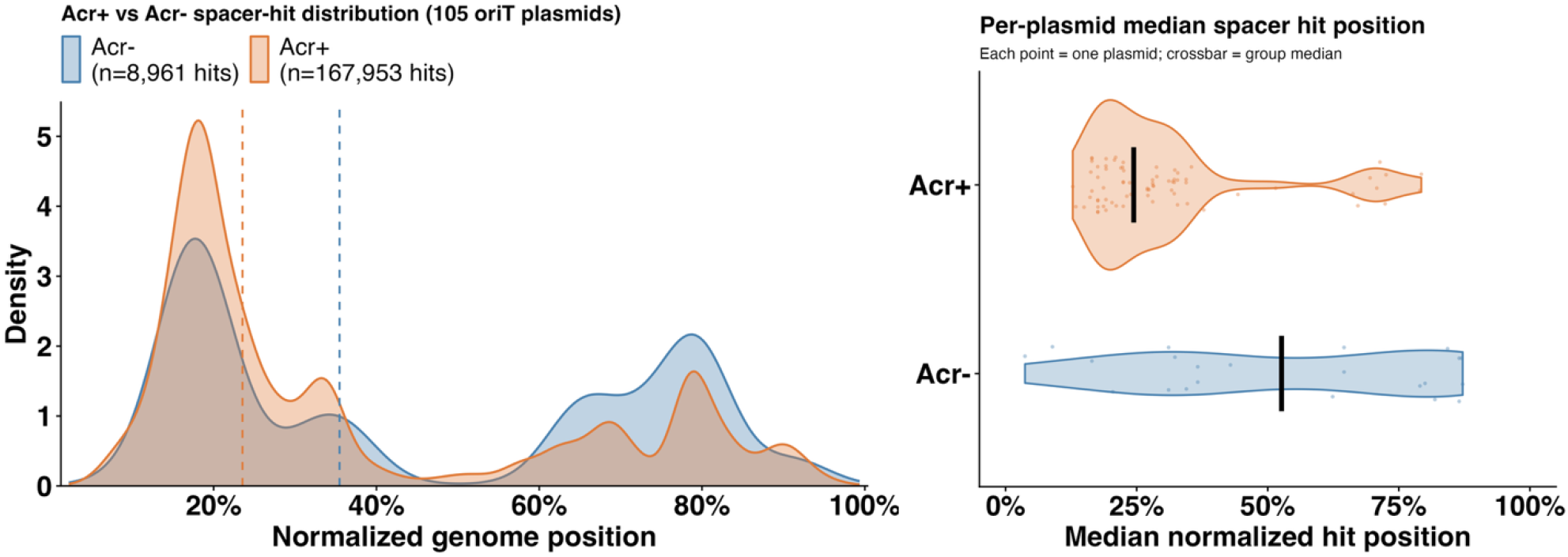
CRISPR spacer hits occur earlier in plasmid genomes carrying anti-CRISPR systems. Plasmids were classified as Acr+ or Acr-. Genome positions were oriented relative to the relaxase and *oriT*, with normalized position 0 representing the leading region that enters the recipient cell first during conjugative transfer. **Left)** Density distributions of 176,914 spacer-hit positions across normalized plasmid genomes, separated by Acr status. Dashed lines indicate the median spacer-hit position for each group. **Right)** Spacer-hit positions were summarized within each plasmid by their median normalized position, such that each point represents one plasmid. Black crossbars indicate the median of these per-plasmid values within each group. Median spacer-hit positions occurred earlier in Acr+ plasmids than in Acr- plasmids (23.5% versus 35.5%; Hodges–Lehmann shift = −0.175, p = 1.95 × 10□ □).

### Limitations

Our datasets are incomplete in two crucial ways. First, there is probably a bias for the identification of Acr defense systems due to recent interest in these systems leading them to be overrepresented. Second, we do not have a complete record of all phages or plasmids. Of the 16,899 spacers queried, 44.8% did not map to any genome in the original phage or plasmid BLAST databases, and over half of all phages did not have any spacer hits. This suggests a wider diversity of phages and plasmids that are yet to be identified.

Analysis of spacers is a form of survivorship bias. For a MGE’s nucleic acid to be in a CRISPR array, the host that acquired the spacer must have survived and replicated. In this way we could expect that spacers represent the less fit members from a MGE population. On the other hand, there is likely a selective advantage to the host for spacers to target the most ‘virulent’ phages and plasmids. It remains to be seen how these opposing forces balance out.

As a final note of interest, as described above (Figure 3C) we demonstrate that spacers can map to regions outside of the expected density of certain genes. Due to this expansion, we advance several different hypotheses. First, building on the Phage Time Continuum, genes that are “out of place” reduce the fitness of phages, leading to them being captured more often in spacers, which we see as a survivorship bias. Additionally, we notice that the genes of the anti-defense system targeted are typically the first to enter the cell. For example, in phages, there is a clear positional displacement of spacers mapping to regions outside of the expected density of the anti-RM systems (Figure 2C). This potentially represents the first anti-RM genes encountered during infection; however, it also represents a distinct RM gene product: the *mom* (phage Mu DNA-modification protein) [40, 41]. In the two aforementioned cases, The Phage Time Continuum directly impacts fitness of the phage where gene position and thus timing manipulates the likelihood of CRISPR-Cas being successful, we will explore this in a subsequent publication.

### Conclusion

CRISPR does not randomly target phage or plasmid genes; we show both the function and position of the gene is important. In a possible example of selfish behavior, the CRISPR-Cas system of *P. aeruginosa* preferentially captures spacers that match back to anti-CRISPR genes over other anti-defense systems. When the MGE can defend itself against CRISPR-Cas, spacers are taken from earlier positions. These data point to the idea that CRISPR-Cas applies selection on MGEs across multiple levels. First, phages and plasmids are under selection for both the specific function of a gene, as shown by the increased abundance of Acr systems in spacers. Second, the overall architecture of their genomes is under selection, as demonstrated by the shift in spacer position and by the data showing an enrichment of spacers targeting anti-defense systems when they fall outside the expected position in phages.

## Methods

### Genome Datasets and Pre-processing

A total of 1,770□*Pseudomonas aeruginosa*□bacteriophage genomes and 450□*P. aeruginosa*□plasmid genomes were used as the reference sequence sets. Phage genomes greater than 100 kb were excluded from positional analyses, as jumbo phages have fundamentally different genomic architectures and expression programs [42-44]. While jumbo phages are typically defined as 200 kb or larger, preliminary taxonomic and architectural analyses of the 37 phages between 100-200 kb were unable to cleanly distinguish jumbo from non-jumbo genomes in this intermediate range. We compared taxnomy and found no clear signature of jumbo phage in the intermediate group. We also compared the genomic organization (number of geness and directionalility of operons) and again found no clear concencise to distinguish these phage from jumbo phages. The 100 kb threshold was therefore retained as a conservative cutoff (SI table 6 & 7). Phage genomes were dereplicated at 95% average nucleotide identity (ANI), requiring 95% query coverage, such that genomes meeting both thresholds were collapsed into a single representative that could be mapped back to the original genomes. Phages absent from the dereplication mapping were retained as singleton representatives. After dereplication and filtering to genomes 5-100 kb in length, 1,462 phage genomes remained. No size filter was applied to plasmids, as there is not an equivalent size-based distinction between plasmids analogous to those distinguishing jumbo from non-jumbo phages[19].

### Genome Annotation

Phage genomes were annotated using Pharokka v1.7.5 with the August 2023 PHROGs database and Phanotate as the gene predictor [45-47], producing gene coordinates, functional categories, and annotation labels for each coding sequence.

### Genome Reorientation

All phage and plasmid genomes were reoriented to a consistent reference frame prior to positional analysis using DNAapler v1.3.0 [48]. Phage genomes were oriented using the large terminase subunit gene (*terL*) as the reference, placing *terL* near genome position 0. Plasmid genomes were oriented following the leading-region framework described by Samuel *et al*. (2024) [18]. Mobilizable and conjugative plasmids were identified based on the presence of conjugation-associated features, and relaxase proteins were detected by hmmsearch (HMMER v3.4). DNAapler was run in custom mode using the MOB-suite (v3.1.9) relaxase protein database to place the relaxase near position 0. The origin of transfer (*oriT*) was identified independently by nucleotide BLAST (v2.17.0) against the MOB-suite *oriT* database. Plasmids were required to have a relaxase-*oriT* distance of no more than 3,500 bp, following Samuel *et al*.; 105 plasmids met this criterion and were retained for positional analyses.

Because DNAapler initially linearized plasmid genomes with the relaxase near position 0, *oriT* occurred near the opposite end of the linear representation as a consequence of their adjacency on the circular plasmid. The best *oriT* hit occurred at a median normalized position of 99.1%, consistent with *oriT* lying immediately upstream of the relaxase. Plasmid coordinates were therefore inverted using (1-x), such that normalized position 0 corresponded to *oriT* and the beginning of the leading region transferred into the recipient first, while position 1 corresponded to the relaxase and lagging end of the transferred genome.

Phage orientation was evaluated by examining the position of TerL in the reoriented genomes. In the original 1,770-genome dataset, TerL was recovered in 1,680 genomes (94.9%), with a median normalized position of 1.33% and 97.4% of detected TerL proteins occurring within the first 5% of the genome. For plasmids, orientation of the final 105-genome dataset was evaluated by the position of the relaxase in the reoriented genome and its proximity to *oriT*. Relaxases occurred at a median normalized position of 2.46%, with 96.2% within the first 5% of the pre-flip genome, while all retained plasmids contained an *oriT* within 3,500 bp of the relaxase.

### Genome Length Normalization

Genome length was calculated from the total nucleotide length of each reoriented FASTA sequence. Feature positions were represented by their midpoint coordinates. Gene midpoints were calculated as ((start + stop)/2), while spacer-hit midpoints were calculated as ((min(sstart, send) + max(sstart, send))/2) to account for reverse-strand alignments. Midpoint coordinates were divided by total genome length to produce normalized positions on a [0,1] scale. For plasmids, normalized coordinates were subsequently inverted as (1-x), as described above, so that position 0 corresponded to the beginning of the leading region at *oriT*. Phage position 0 corresponded to the TerL-oriented genome start.

### Identification of Anti-Defense Systems

Anti-defense systems were annotated in each reoriented genome using DefenseFinder’s --antidefensefinder-only option[49-51]. DefenseFinder v2.0.1 (models v2.0.2) was run on the reoriented protein sequences (.prt files) for each genome. System annotations include the system type (e.g., Anti-CRISPR, Anti-RM), subtype, and the nucleotide coordinates of each system within the reoriented genome. Genomic positions of anti-defense systems were normalized to a [0, 1] scale by dividing the system midpoint nucleotide coordinate by total genome length. The system midpoint was computed as the mean of the system’s start and end nucleotide coordinates.

### *P. aeruginosa*□CRISPR Spacer Database

*P. aeruginosa*□CRISPR spacer sequences were obtained from Ortega-Sanz *et al*., 2023 [52], a curated collection of spacers extracted from the CRISPR arrays of sequenced *P. aeruginosa isolates*. These spacers represent the recorded immune history of□*P. aeruginosa*□CRISPR-Cas systems and serve as the query sequences for all alignment analyses. Each spacer is a fragment of a mobile genetic element (phage or plasmid) that was previously acquired into a CRISPR array and therefore marks a past infection or invasion event. The dataset comprised 184,972 spacer sequences, which served as the query sequences for all alignment analyses. Because a spacer’s position within a target genome reflects where CRISPR-Cas sampled that element, the full set provides a genome-wide record of which regions of phages and plasmids have historically been targeted by *P. aeruginosa* immunity.

### Alignment of Spacers to Phage and Plasmid Genomes

*P. aeruginosa* CRISPR spacers were aligned to all reoriented phage and plasmid genomes using BLASTN (v2.17.0) [53, 54]. Spacers were aligned separately against the 1,462 reoriented phage genomes and the 105 *oriT*- anchored plasmid genomes, using identical alignment parameters for both sequence sets. In each case, BLASTN was run with the blastn-short task, requiring 95% percent identity (-perc_identity 95) and full-length spacer coverage (-qcov_hsp_perc 100), such that only high-confidence, full-coverage spacer matches were retained. The resulting BLAST output (tabular format 6) provides, for each spacer hit, the query spacer identity, the subject genome accession, the alignment start and end coordinates within the subject genome (sstart, send), and alignment statistics.

### Identification of Spacer Hits Overlapping Anti-Defense Systems

To identify spacer hits that physically overlap annotated anti-defense system coordinates, genomic interval overlap analysis was performed between the BLAST hit coordinates and the DefenseFinder system coordinates for each genome. Overlapping spacer hits were retained in a separate overlaps dataset that records both the hit position and the type and subtype of the anti-defense system that the spacer targets. This dataset was used for all analyses of anti-defense system targeting rates and for the positional distribution comparisons between anti-defense system locations and the genomic positions where spacers hit them.

### Anti-Defense Targeting Enrichment Analysis

For each anti-defense class, targeting intensity was calculated as mean spacer hits per annotated system. Acr systems were compared with all other anti-defense systems combined, separately for phages and plasmids. Because spacer hits were non-independent across genomes and systems, these comparisons were treated descriptively. Raw system and spacer-hit counts were also reported.

### Positional Distribution Modeling

To characterize the genomic position distributions of anti-defense systems and their spacer hit locations, Gaussian mixture models were fitted independently to three distributions for each of the four dataset-by-type combinations (phage Anti-CRISPR, phage Anti-RM, plasmid Anti-CRISPR, plasmid Anti-RM):

**Distribution A:** □The normalized midpoint positions of all annotated anti-defense systems of that type, representing where anti-defense genes are located in the genome. **Distribution B1:** □The normalized midpoint positions of spacer hits overlapping those systems, representing where in the genome spacers physically land on anti-defense loci. **Distribution B2:** □The normalized midpoint positions of the□systems□that are hit by spacers, representing which systems (by genomic position) are targeted, regardless of where within the system the spacer lands.

Gaussian mixture modeling was performed using the□mclust v6.0.1□package in R, with the number of mixture components selected by Bayesian Information Criterion (BIC)[55]. Kolmogorov–Smirnov (KS) tests were used to assess whether distributions A, B1, and B2 differed significantly from one another for each panel.

Positional distributions were additionally visualized as mirrored kernel density plots with anti-defense system positions shown on the upper half of the y-axis and spacer hit positions reflected on the lower half, facilitating direct visual comparison of the two distributions at each genomic position.

### Acr+ vs Acr- Spacer Positional Analysis

Phage and plasmid genomes were classified as Acr+ if they contained at least one Acr system detected by DefenseFinder and as Acr- if no Acr system was detected. Following dereplication and genome-length filtering, the phage dataset contained 1,462 genomes, including 98 Acr+ and 1,364 Acr- phages. The plasmid analysis was restricted to the final set of 105 mobilizable or conjugative plasmids with validated *oriT*-relaxase organization, including 76 Acr+ and 29 Acr- plasmids.

All retained spacer BLAST hits were included in the positional analysis, rather than restricting the dataset to spacer hits overlapping anti-defense systems. Normalized spacer-hit midpoint positions were calculated as described above. Three complementary comparisons were performed for both phages and plasmids.

First, the proportion of genomes in each Acr group targeted by at least one spacer was compared using Fisher’s exact test. Second, normalized spacer-hit positions were pooled within each Acr group and compared using a one-sided Wilcoxon rank-sum test, with the alternative hypothesis that spacer hits occurred earlier in Acr+ genomes than in Acr- genomes. Hodges-Lehmann estimates were used to quantify the positional shift between groups.

Third, to account for differences in the number of spacer hits received by individual genomes, the median normalized spacer-hit position was calculated independently for each genome with at least one spacer hit. These per-genome median positions were then compared between Acr+ and Acr- groups using a one-sided Wilcoxon rank-sum test with Hodges-Lehmann estimation. This approach gives each targeted genome equal weight regardless of the number of spacer hits it received.

All retained spacer BLAST hits were included in the primary positional analysis, including hits that directly overlapped Acr loci. To determine whether the observed Acr+ versus Acr- positional shift was driven by direct targeting of Acr systems, a separate sensitivity analysis was performed in which spacer hits overlapping annotated Anti-CRISPR loci were removed from Acr+ genomes. The pooled hit-level and per-genome median comparisons were then repeated. Excluding Acr-locus hits produced no meaningful change in the observed positional pattern, so the full dataset was retained for the primary analyses and figures (SI Figs X & X).

## Statistical Analyses

All statistical analyses were performed in R version 4.2.2 [56]. Group comparisons of spacer-hit positions used one-sided Wilcoxon rank-sum tests with Hodges–Lehmann estimation of the positional shift; proportions of targeted genomes were compared using Fisher’s exact test; and distributional differences were assessed using Kolmogorov–Smirnov tests. Gaussian mixture modeling was performed using mclust v6.0.1, with the number of components selected by Bayesian Information Criterion. Statistical details for each analysis are given in the corresponding subsections above.

## Notes

### Competing Interest Statement

The authors have declared no competing interest.

### Funding

- NIH: r35gm166251

