## Supplementary material for "The Phage Time Continuum: Genome Organization Constrains Anti-CRISPR Success": SI figures and tables

**Supplementary information.**

Sean Czerwinski^a^, Iris Irby ^b^, Gabriel Morales Artavia^a^, Sam P Brown ^b^, James Gurney^aφ^

^a^ School of Biology, Georgia State University, Atlanta, Georgia, USA; ^b^ Center for Microbial Dynamics & Infection, School of Biological Sciences, Georgia Institute of Technology, Atlanta, Georgia, USA.

SI Table 1. Anti-defense targeting intensity by system class, for phages and plasmids. For each anti-defense class, the number of annotated systems (n_systems), the number of overlapping spacer hits (n_hits), and the mean spacer hits per system (hits_per_system) are reported separately for phage and plasmid genomes. Acr systems are the most intensively targeted class in both mobile-element types (42.7 hits/system in phages; 4.2 in plasmids), substantially exceeding all other anti-defense classes.

| type_display | mge_type | n_systems | n_hits | hits_per_system |
| --- | --- | --- | --- | --- |
| Acr | Phage | 158 | 6740 | 42.6582278 |
| Acr | Plasmid | 204 | 851 | 4.17156863 |
| Anti-CBASS | Phage | 33 | 304 | 9.21212121 |
| Anti-CBASS | Plasmid | 2 | 0 | 0 |
| Anti-Dnd | Phage | 19 | 12 | 0.63157895 |
| Anti-RM | Phage | 60 | 106 | 1.76666667 |
| Anti-RM | Plasmid | 102 | 55 | 0.53921569 |
| Anti-Retron | Phage | 2 | 0 | 0 |
| Anti-Thoeris | Phage | 2 | 0 | 0 |
| Mok/Hok/Sok | Plasmid | 1 | 0 | 0 |
| NADP-related | Phage | 122 | 0 | 0 |
| Other | Phage | 2 | 0 | 0 |

SI Table 2. Per-genus spacer-targeting summary for classified phage genera. For each genus with assigned taxonomy, the number of phages (n_phages), the percentage receiving zero spacer hits (pct_0_hits), the percentage targeted by ≥1 spacer (pct_targeted), and the median, mean, length-normalized median (median_hits_per_kb), maximum, and total spacer hits are reported. Genera are included where taxonomic assignment was available; targeting intensity varies markedly across genera.

| genus | n_phages | pct_0_hits | pct_targeted | median_hits | mean_hits | median_hits_per_kb | max_hits | total_hits |
| --- | --- | --- | --- | --- | --- | --- | --- | --- |
| Casadabanvirus | 64 | 0 | 100 | 6729 | 6343.2 | 194.31 | 9541 | 405965 |
| Mechnikovvirus | 6 | 0 | 100 | 3585.5 | 3616.3 | 92.16 | 4110 | 21698 |
| Citexvirus | 5 | 0 | 100 | 3329 | 2996.8 | 90.11 | 3492 | 14984 |
| Beetrevirus | 6 | 0 | 100 | 3282.5 | 2968.7 | 84.76 | 3560 | 17812 |
| Detrevirus | 8 | 0 | 100 | 2902.5 | 2951.1 | 55.6 | 5400 | 23609 |
| Torontovirus | 6 | 0 | 100 | 2155.5 | 2236.2 | 56.2 | 2944 | 13417 |
| Hollowayvirus | 20 | 0 | 100 | 1644.5 | 1652.6 | 26.6 | 2450 | 33052 |
| Pbunavirus | 195 | 0 | 100 | 31 | 102.9 | 0.48 | 529 | 20064 |
| Paundecimvirus | 9 | 0 | 100 | 25 | 204.1 | 0.5 | 446 | 1837 |
| Samunavirus | 11 | 0 | 100 | 4 | 40.5 | 0.05 | 208 | 445 |
| Nankokuvirus | 30 | 16.7 | 83.3 | 14 | 53.1 | 0.16 | 428 | 1592 |
| Kochitakasuvirus | 4 | 25 | 75 | 209 | 239.5 | 3.29 | 540 | 958 |
| Septimatrevirus | 41 | 43.9 | 56.1 | 2 | 1.1 | 0.05 | 2 | 46 |
| Tunggulvirus | 3 | 66.7 | 33.3 | 0 | 1419 | 0 | 4257 | 4257 |
| Bruynoghevirus | 70 | 71.4 | 28.6 | 0 | 0.3 | 0 | 1 | 20 |
| Pakpunavirus | 73 | 94.5 | 5.5 | 0 | 2.4 | 0 | 170 | 173 |
| Abidjanvirus | 8 | 100 | 0 | 0 | 0 | 0 | 0 | 0 |
| Bifseptvirus | 6 | 100 | 0 | 0 | 0 | 0 | 0 | 0 |
| Flaumdravirus | 5 | 100 | 0 | 0 | 0 | 0 | 0 | 0 |
| Ghunavirus | 42 | 100 | 0 | 0 | 0 | 0 | 0 | 0 |
| Hennigervirus | 7 | 100 | 0 | 0 | 0 | 0 | 0 | 0 |
| Iggyvirus | 5 | 100 | 0 | 0 | 0 | 0 | 0 | 0 |
| Jamesmcgillvirus | 9 | 100 | 0 | 0 | 0 | 0 | 0 | 0 |
| Kremarvirus | 3 | 100 | 0 | 0 | 0 | 0 | 0 | 0 |
| Litunavirus | 45 | 100 | 0 | 0 | 0 | 0 | 0 | 0 |
| Luzseptimavirus | 9 | 100 | 0 | 0 | 0 | 0 | 0 | 0 |
| Nerthusvirus | 5 | 100 | 0 | 0 | 0 | 0 | 0 | 0 |
| Nipunavirus | 9 | 100 | 0 | 0 | 0 | 0 | 0 | 0 |
| Otagovirus | 15 | 100 | 0 | 0 | 0 | 0 | 0 | 0 |
| Pamexvirus | 5 | 100 | 0 | 0 | 0 | 0 | 0 | 0 |
| Pfluvirus | 4 | 100 | 0 | 0 | 0 | 0 | 0 | 0 |
| Phikmvvirus | 64 | 100 | 0 | 0 | 0 | 0 | 0 | 0 |
| Pifdecavirus | 24 | 100 | 0 | 0 | 0 | 0 | 0 | 0 |
| Pijolavirus | 3 | 100 | 0 | 0 | 0 | 0 | 0 | 0 |
| Pollyceevirus | 3 | 100 | 0 | 0 | 0 | 0 | 0 | 0 |
| Readingvirus | 3 | 100 | 0 | 0 | 0 | 0 | 0 | 0 |
| Tartuvirus | 4 | 100 | 0 | 0 | 0 | 0 | 0 | 0 |
| Uliginvirus | 3 | 100 | 0 | 0 | 0 | 0 | 0 | 0 |
| Unosvirus | 6 | 100 | 0 | 0 | 0 | 0 | 0 | 0 |
| Warsawvirus | 3 | 100 | 0 | 0 | 0 | 0 | 0 | 0 |
| Yuavirus | 43 | 100 | 0 | 0 | 0 | 0 | 0 | 0 |
| Zurivirus | 3 | 100 | 0 | 0 | 0 | 0 | 0 | 0 |

SI Table 3. Pooled hit-level Wilcoxon rank-sum test comparing normalized spacer-hit positions between Acr+ and Acr− phages. One-sided test (alternative: Acr+ earlier than Acr−), with the Hodges–Lehmann estimate of the positional shift and the number of hits per group. The large sample size (n = 583,709 Acr+ vs 453,938 Acr− hits) yields a vanishingly small p-value; the Hodges–Lehmann shift (−0.049) is the more informative effect-size measure.

| test | W_statistic | p_value | HL_estimate | n_acr_plus | n_acr_minus |
| --- | --- | --- | --- | --- | --- |
| Wilcoxon rank-sum (one-sided: Acr+ < Acr-) | 1.1849E+11 | 0 | -0.0490901 | 583709 | 453938 |

SI Table 4. Phage positional-shift statistics, pooled and per-genome. Results of two one-sided Wilcoxon rank-sum tests (alternative: Acr+ earlier): the pooled hit-level test (all spacer hits) and the per-phage median test (each genome contributing a single median position). The per-phage test (Hodges–Lehmann = -0.128, p = 4.2 × 10⁻⁸; n = 98 Acr+, 574 Acr-) weights each genome equally and is robust to the small number of heavily targeted phages.

| test | W | p_value | HL_estimate | n_pos | n_neg |
| --- | --- | --- | --- | --- | --- |
| Pooled hit-position Wilcoxon (Acr-locus excluded, Acr+<Acr-) | 1.1818E+11 | 0 | -0.0467058 | 578983 | 453938 |
| Per-phage median Wilcoxon (Acr-locus excluded, Acr+<Acr-) | 18605 | 4.16E-08 | -0.127618 | 98 | 574 |

SI Table 5 Plasmid positional-shift statistics, pooled and per-genome. Results of two one-sided Wilcoxon rank-sum tests comparing normalized spacer-hit positions between Acr+ and Acr- plasmids. The pooled test (Hodges–Lehmann = -0.027, p = 1.2 × 10⁻³) and the per-plasmid median test (Hodges–Lehmann = -0.184, p = 2.4 × 10⁻⁴; n = 76 Acr+, 22 Acr−) are reported.

| test | W | p_value | HL_estimate | n_pos | n_neg |
| --- | --- | --- | --- | --- | --- |
| Pooled hit-position Wilcoxon (Acr-locus excluded, Acr+<Acr-) | 1681578 | 0.00117595 | -0.0269568 | 8112 | 453 |
| Per-plasmid median Wilcoxon (Acr-locus excluded, Acr+<Acr-) | 426 | 2.44E-04 | -0.1840312 | 76 | 22 |

SI Table 6.Taxonomic classification of the 37 phage genomes in the 100–200 kb intermediate size range. NCBI taxonomy (taxid, organism, family, genus) for each phage excluded from positional analyses by the 100 kb threshold. Eight phages belong to recognized large-genome families (Straboviridae, Ackermannviridae, Phycodnaviridae); the remainder are predominantly Pseudomonas-infecting genera not clearly classifiable as jumbo.

| accession | taxid | organism | family | genus |
| --- | --- | --- | --- | --- |
| AF503408 | 10682 | Enterobacteria phage P7 | NA | Punavirus |
| AY375531 | 2907963 | Aeromonas phage 44RR2.8t | Straboviridae | Biquartavirus |
| AY962392 | 321023 | Aeromonas phage 31 | Straboviridae | Biquartavirus |
| EF469154 | 293178 | Escherichia phage JS98 | Straboviridae | Dhakavirus |
| JN225873 | 1120767 | Ostreococcus tauri virus RT-2011 | Phycodnaviridae | Prasinovirus |
| KU862660 | 1815582 | Pseudomonas phage phiPMW | NA | Plaisancevirus |
| KU873925 | 1815630 | Pseudomonas phage pf16 | NA | Chakrabartyvirus |
| KY971610 | 1983556 | Pseudomonas phage PspYZU05 | Straboviridae | Jiangsuvirus |
| MG018927 | 2048977 | Pseudomonas phage nickie | NA | Nickievirus |
| MK813938 | 2588515 | Aeromonas phage 2L372X | NA | Plateaulakevirus |
| MN871475 | 2698992 | Pseudomonas phage Pa-sanjiao | Ackermannviridae | Kuttervirus |
| MT354569 | 2755401 | Pseudomonas phage phiK7B1 | NA | Nickievirus |
| MT670417 | 1460361 | Pseudomonas phage phiPsa267 | Vandenendeviridae | Otagovirus |
| MZ605293 | 2856997 | Pseudomonas phage UAVern | Vandenendeviridae | Uavernvirus |
| OM140839 | 2909584 | Pseudomonas phage ZY21 | NA | Nickievirus |
| OM223115 | 2912595 | Pseudomonas phage hairong | NA | Nickievirus |
| OM982621 | 2928830 | Pseudomonas phage Astolliot | Vandenendeviridae | Otagovirus |
| OQ183418 | 3020390 | Pseudomonas phage vB_PpS_SYP | NA | NA |
| OQ992558 | 3046821 | Escherichia phage L14 | Straboviridae | Tequatrovirus |
| OR584013 | 3088885 | Pseudomonas phage PsgM02F | NA | NA |
| OR584014 | 3088886 | Pseudomonas phage PsgM04F | NA | NA |
| OR885871 | 3116514 | Escherichia phage vB_EcoM_JN03 | Ackermannviridae | Kuttervirus |
| PP179310 | 3138546 | Pseudomonas phage Pyxpy01 | NA | NA |
| PP179318 | 3138540 | Pseudomonas phage Nican01 | NA | Nickievirus |
| PP179332 | 3138529 | Pseudomonas phage Cygsa01 | NA | NA |
| PP496437 | 3132641 | Pseudomonas phage vB_PpuM-Aura | NA | NA |
| PP596839 | 3136767 | Pseudomonas phage WP1 | NA | NA |
| PQ464597 | 3374953 | Pseudomonas phage Milchi | NA | NA |
| PQ516283 | 3388586 | Pseudomonas phage vB_PQ1 | NA | NA |
| PQ859842 | 3399671 | Pseudomonas phage CP-p-PA-21011 | NA | NA |
| PV055685 | 3401910 | Pseudomonas phage QdP_Psa3 | NA | NA |
| PV446632 | 3414728 | Pseudomonas phage vB_Pa-PAC11 | NA | NA |
| PV654520 | 3425914 | Pseudomonas phage vB_PSGBLB0006 | NA | NA |
| PV714129 | 3434943 | Escherichia phage Mizar | NA | NA |
| PX583109 | 3473746 | Pseudomonas phage CYPPA010 | NA | NA |
| PX866597 | 3591852 | Bacillus phage Asciepus | NA | NA |
| PX866601 | 3591853 | Bacillus phage Nierus | NA | NA |

SI Table 7. Co-directional gene-run (directon) architecture across phage size bands. For each phage in the 100–200 kb intermediate range and a reference set of ≤100 kb analysis-set genomes, the number of predicted genes (n_genes), the number of maximal same-strand gene runs (n_directons), and the mean genes per directon (genes_per_directon) are reported. Phages are grouped into a reference set (≤100 kb analysis-set sample, n=200), the 100–120 kb band (n=14), a 120–150 kb grey zone (n=2), 150–200 kb phages lacking a recognized large-phage family assignment (n=13), and phages in recognized large-genome families (Straboviridae, Ackermannviridae, Phycodnaviridae; n=8) but fall below the 200 kb threshold for jumbo designation. Direction density does not cleanly separate the 100–120 kb band from the sub-100 kb analysis set, supporting the use of a conservative 100 kb cutoff.

| accession | group | tsv | n_genes | n_directons | genes_per_directon |
| --- | --- | --- | --- | --- | --- |
| AF503408 | notjumbo_100_120 | /Volumes/Extreme_SSD/CRISPR_spacer_analyses/Paper3/pharokka_results/AF503408.1/AF503408.1_cds_final_merged_output.tsv | 148 | 35 | 4.22857143 |
| AY375531 | jumbo_family | /Volumes/Extreme_SSD/CRISPR_spacer_analyses/Paper3/pharokka_results/AY375531.1/AY375531.1_cds_final_merged_output.tsv | 277 | 23 | 12.0434783 |
| AY962392 | jumbo_family | /Volumes/Extreme_SSD/CRISPR_spacer_analyses/Paper3/pharokka_results/AY962392.1/AY962392.1_cds_final_merged_output.tsv | 278 | 27 | 10.2962963 |
| EF469154 | jumbo_family | /Volumes/Extreme_SSD/CRISPR_spacer_analyses/Paper3/pharokka_results/EF469154.1/EF469154.1_cds_final_merged_output.tsv | 297 | 13 | 22.8461538 |
| JN225873 | jumbo_family | /Volumes/Extreme_SSD/CRISPR_spacer_analyses/Paper3/pharokka_results/JN225873.1/JN225873.1_cds_final_merged_output.tsv | 285 | 103 | 2.76699029 |
| KU862660 | notjumbo_100_120 | /Volumes/Extreme_SSD/CRISPR_spacer_analyses/Paper3/pharokka_results/KU862660/KU862660_cds_final_merged_output.tsv | 246 | 13 | 18.9230769 |
| KU873925 | large_unresolved_150_200 | /Volumes/Extreme_SSD/CRISPR_spacer_analyses/Paper3/pharokka_results/KU873925/KU873925_cds_final_merged_output.tsv | 260 | 19 | 13.6842105 |
| KY971610 | jumbo_family | /Volumes/Extreme_SSD/CRISPR_spacer_analyses/Paper3/pharokka_results/KY971610/KY971610_cds_final_merged_output.tsv | 234 | 19 | 12.3157895 |
| MG018927 | notjumbo_100_120 | /Volumes/Extreme_SSD/CRISPR_spacer_analyses/Paper3/pharokka_results/MG018927/MG018927_cds_final_merged_output.tsv | 177 | 13 | 13.6153846 |
| MK813938 | notjumbo_100_120 | /Volumes/Extreme_SSD/CRISPR_spacer_analyses/Paper3/pharokka_results/MK813938.1/MK813938.1_cds_final_merged_output.tsv | 235 | 24 | 9.79166667 |
| MN871475 | jumbo_family | /Volumes/Extreme_SSD/CRISPR_spacer_analyses/Paper3/pharokka_results/MN871475/MN871475_cds_final_merged_output.tsv | 237 | 29 | 8.17241379 |
| MT354569 | notjumbo_100_120 | /Volumes/Extreme_SSD/CRISPR_spacer_analyses/Paper3/pharokka_results/MT354569/MT354569_cds_final_merged_output.tsv | 182 | 7 | 26 |
| MT670417 | notjumbo_100_120 | /Volumes/Extreme_SSD/CRISPR_spacer_analyses/Paper3/pharokka_results/MT670417/MT670417_cds_final_merged_output.tsv | 204 | 9 | 22.6666667 |
| MZ605293 | notjumbo_100_120 | /Volumes/Extreme_SSD/CRISPR_spacer_analyses/Paper3/pharokka_results/MZ605293/MZ605293_cds_final_merged_output.tsv | 216 | 12 | 18 |
| OM140839 | notjumbo_100_120 | /Volumes/Extreme_SSD/CRISPR_spacer_analyses/Paper3/pharokka_results/OM140839/OM140839_cds_final_merged_output.tsv | 186 | 9 | 20.6666667 |
| OM223115 | notjumbo_100_120 | /Volumes/Extreme_SSD/CRISPR_spacer_analyses/Paper3/pharokka_results/OM223115/OM223115_cds_final_merged_output.tsv | 192 | 11 | 17.4545455 |
| OM982621 | large_unresolved_150_200 | /Volumes/Extreme_SSD/CRISPR_spacer_analyses/Paper3/pharokka_results/OM982621/OM982621_cds_final_merged_output.tsv | 327 | 41 | 7.97560976 |
| OQ183418 | notjumbo_100_120 | /Volumes/Extreme_SSD/CRISPR_spacer_analyses/Paper3/pharokka_results/OQ183418/OQ183418_cds_final_merged_output.tsv | 251 | 11 | 22.8181818 |
| OQ992558 | jumbo_family | /Volumes/Extreme_SSD/CRISPR_spacer_analyses/Paper3/pharokka_results/OQ992558.1/OQ992558.1_cds_final_merged_output.tsv | 286 | 19 | 15.0526316 |
| OR584013 | large_unresolved_150_200 | /Volumes/Extreme_SSD/CRISPR_spacer_analyses/Paper3/pharokka_results/OR584013/OR584013_cds_final_merged_output.tsv | 278 | 13 | 21.3846154 |
| OR584014 | large_unresolved_150_200 | /Volumes/Extreme_SSD/CRISPR_spacer_analyses/Paper3/pharokka_results/OR584014/OR584014_cds_final_merged_output.tsv | 324 | 17 | 19.0588235 |
| OR885871 | jumbo_family | /Volumes/Extreme_SSD/CRISPR_spacer_analyses/Paper3/pharokka_results/OR885871.1/OR885871.1_cds_final_merged_output.tsv | 237 | 23 | 10.3043478 |
| PP179310 | notjumbo_100_120 | /Volumes/Extreme_SSD/CRISPR_spacer_analyses/Paper3/pharokka_results/PP179310/PP179310_cds_final_merged_output.tsv | 213 | 17 | 12.5294118 |
| PP179318 | notjumbo_100_120 | /Volumes/Extreme_SSD/CRISPR_spacer_analyses/Paper3/pharokka_results/PP179318/PP179318_cds_final_merged_output.tsv | 186 | 11 | 16.9090909 |
| PP179332 | large_unresolved_150_200 | /Volumes/Extreme_SSD/CRISPR_spacer_analyses/Paper3/pharokka_results/PP179332/PP179332_cds_final_merged_output.tsv | 286 | 29 | 9.86206897 |
| PP496437 | large_unresolved_150_200 | /Volumes/Extreme_SSD/CRISPR_spacer_analyses/Paper3/pharokka_results/PP496437/PP496437_cds_final_merged_output.tsv | 255 | 15 | 17 |
| PP596839 | notjumbo_100_120 | /Volumes/Extreme_SSD/CRISPR_spacer_analyses/Paper3/pharokka_results/PP596839/PP596839_cds_final_merged_output.tsv | 323 | 38 | 8.5 |
| PQ464597 | large_unresolved_150_200 | /Volumes/Extreme_SSD/CRISPR_spacer_analyses/Paper3/pharokka_results/PQ464597/PQ464597_cds_final_merged_output.tsv | 328 | 29 | 11.3103448 |
| PQ516283 | large_unresolved_150_200 | /Volumes/Extreme_SSD/CRISPR_spacer_analyses/Paper3/pharokka_results/PQ516283/PQ516283_cds_final_merged_output.tsv | 467 | 43 | 10.8604651 |
| PQ859842 | grey_120_150 | /Volumes/Extreme_SSD/CRISPR_spacer_analyses/Paper3/pharokka_results/PQ859842/PQ859842_cds_final_merged_output.tsv | 195 | 9 | 21.6666667 |
| PV055685 | notjumbo_100_120 | /Volumes/Extreme_SSD/CRISPR_spacer_analyses/Paper3/pharokka_results/PV055685/PV055685_cds_final_merged_output.tsv | 169 | 15 | 11.2666667 |
| PV446632 | large_unresolved_150_200 | /Volumes/Extreme_SSD/CRISPR_spacer_analyses/Paper3/pharokka_results/PV446632.1/PV446632.1_cds_final_merged_output.tsv | 337 | 33 | 10.2121212 |
| PV654520 | grey_120_150 | /Volumes/Extreme_SSD/CRISPR_spacer_analyses/Paper3/pharokka_results/PV654520/PV654520_cds_final_merged_output.tsv | 237 | 15 | 15.8 |
| PV714129 | large_unresolved_150_200 | /Volumes/Extreme_SSD/CRISPR_spacer_analyses/Paper3/pharokka_results/PV714129.1/PV714129.1_cds_final_merged_output.tsv | 280 | 19 | 14.7368421 |
| PX583109 | large_unresolved_150_200 | /Volumes/Extreme_SSD/CRISPR_spacer_analyses/Paper3/pharokka_results/PX583109.1/PX583109.1_cds_final_merged_output.tsv | 281 | 23 | 12.2173913 |
| PX866597 | large_unresolved_150_200 | /Volumes/Extreme_SSD/CRISPR_spacer_analyses/Paper3/pharokka_results/PX866597.1/PX866597.1_cds_final_merged_output.tsv | 286 | 15 | 19.0666667 |
| PX866601 | large_unresolved_150_200 | /Volumes/Extreme_SSD/CRISPR_spacer_analyses/Paper3/pharokka_results/PX866601.1/PX866601.1_cds_final_merged_output.tsv | 288 | 11 | 26.1818182 |
| OQ504960 | ref_le100kb | /Volumes/Extreme_SSD/CRISPR_spacer_analyses/Paper3/pharokka_results/OQ504960/OQ504960_cds_final_merged_output.tsv | 102 | 8 | 12.75 |
| MT119367 | ref_le100kb | /Volumes/Extreme_SSD/CRISPR_spacer_analyses/Paper3/pharokka_results/MT119367/MT119367_cds_final_merged_output.tsv | 105 | 14 | 7.5 |
| JX434032 | ref_le100kb | /Volumes/Extreme_SSD/CRISPR_spacer_analyses/Paper3/pharokka_results/JX434032/JX434032_cds_final_merged_output.tsv | 58 | 4 | 14.5 |
| OP310970 | ref_le100kb | /Volumes/Extreme_SSD/CRISPR_spacer_analyses/Paper3/pharokka_results/OP310970/OP310970_cds_final_merged_output.tsv | 100 | 16 | 6.25 |
| PQ758385 | ref_le100kb | /Volumes/Extreme_SSD/CRISPR_spacer_analyses/Paper3/pharokka_results/PQ758385/PQ758385_cds_final_merged_output.tsv | 412 | 51 | 8.07843137 |
| MK510992 | ref_le100kb | /Volumes/Extreme_SSD/CRISPR_spacer_analyses/Paper3/pharokka_results/MK510992/MK510992_cds_final_merged_output.tsv | 73 | 7 | 10.4285714 |
| LC102730 | ref_le100kb | /Volumes/Extreme_SSD/CRISPR_spacer_analyses/Paper3/pharokka_results/LC102730/LC102730_cds_final_merged_output.tsv | 104 | 16 | 6.5 |
| KU948710 | ref_le100kb | /Volumes/Extreme_SSD/CRISPR_spacer_analyses/Paper3/pharokka_results/KU948710/KU948710_cds_final_merged_output.tsv | 103 | 9 | 11.4444444 |
| PP107937 | ref_le100kb | /Volumes/Extreme_SSD/CRISPR_spacer_analyses/Paper3/pharokka_results/PP107937/PP107937_cds_final_merged_output.tsv | 99 | 11 | 9 |
| PP573285 | ref_le100kb | /Volumes/Extreme_SSD/CRISPR_spacer_analyses/Paper3/pharokka_results/PP573285/PP573285_cds_final_merged_output.tsv | 80 | 7 | 11.4285714 |
| MN871461 | ref_le100kb | /Volumes/Extreme_SSD/CRISPR_spacer_analyses/Paper3/pharokka_results/MN871461/MN871461_cds_final_merged_output.tsv | 104 | 14 | 7.42857143 |
| PP496457 | ref_le100kb | /Volumes/Extreme_SSD/CRISPR_spacer_analyses/Paper3/pharokka_results/PP496457/PP496457_cds_final_merged_output.tsv | 55 | 3 | 18.3333333 |
| PQ571160 | ref_le100kb | /Volumes/Extreme_SSD/CRISPR_spacer_analyses/Paper3/pharokka_results/PQ571160/PQ571160_cds_final_merged_output.tsv | 59 | 4 | 14.75 |
| LN610581 | ref_le100kb | /Volumes/Extreme_SSD/CRISPR_spacer_analyses/Paper3/pharokka_results/LN610581/LN610581_cds_final_merged_output.tsv | 173 | 11 | 15.7272727 |
| PV565042 | ref_le100kb | /Volumes/Extreme_SSD/CRISPR_spacer_analyses/Paper3/pharokka_results/PV565042/PV565042_cds_final_merged_output.tsv | 100 | 11 | 9.09090909 |
| AY576273.1 | ref_le100kb | /Volumes/Extreme_SSD/CRISPR_spacer_analyses/Paper3/pharokka_results/AY576273.1/AY576273.1_cds_final_merged_output.tsv | 95 | 1 | 95 |
| OR621024 | ref_le100kb | /Volumes/Extreme_SSD/CRISPR_spacer_analyses/Paper3/pharokka_results/OR621024/OR621024_cds_final_merged_output.tsv | 24 | 3 | 8 |
| MW175491 | ref_le100kb | /Volumes/Extreme_SSD/CRISPR_spacer_analyses/Paper3/pharokka_results/MW175491/MW175491_cds_final_merged_output.tsv | 199 | 13 | 15.3076923 |
| OM982620 | ref_le100kb | /Volumes/Extreme_SSD/CRISPR_spacer_analyses/Paper3/pharokka_results/OM982620/OM982620_cds_final_merged_output.tsv | 200 | 23 | 8.69565217 |
| MK511013 | ref_le100kb | /Volumes/Extreme_SSD/CRISPR_spacer_analyses/Paper3/pharokka_results/MK511013/MK511013_cds_final_merged_output.tsv | 69 | 9 | 7.66666667 |
| PX583109.1 | ref_le100kb | /Volumes/Extreme_SSD/CRISPR_spacer_analyses/Paper3/pharokka_results/PX583109.1/PX583109.1_cds_final_merged_output.tsv | 281 | 23 | 12.2173913 |
| OM937766 | ref_le100kb | /Volumes/Extreme_SSD/CRISPR_spacer_analyses/Paper3/pharokka_results/OM937766/OM937766_cds_final_merged_output.tsv | 102 | 9 | 11.3333333 |
| MG675557.1 | ref_le100kb | /Volumes/Extreme_SSD/CRISPR_spacer_analyses/Paper3/pharokka_results/MG675557.1/MG675557.1_cds_final_merged_output.tsv | 81 | 15 | 5.4 |
| PV620260 | ref_le100kb | /Volumes/Extreme_SSD/CRISPR_spacer_analyses/Paper3/pharokka_results/PV620260/PV620260_cds_final_merged_output.tsv | 56 | 7 | 8 |
| OP882271 | ref_le100kb | /Volumes/Extreme_SSD/CRISPR_spacer_analyses/Paper3/pharokka_results/OP882271/OP882271_cds_final_merged_output.tsv | 93 | 9 | 10.3333333 |
| MZ826353 | ref_le100kb | /Volumes/Extreme_SSD/CRISPR_spacer_analyses/Paper3/pharokka_results/MZ826353/MZ826353_cds_final_merged_output.tsv | 194 | 29 | 6.68965517 |
| OM141125 | ref_le100kb | /Volumes/Extreme_SSD/CRISPR_spacer_analyses/Paper3/pharokka_results/OM141125/OM141125_cds_final_merged_output.tsv | 101 | 7 | 14.4285714 |
| OP310971 | ref_le100kb | /Volumes/Extreme_SSD/CRISPR_spacer_analyses/Paper3/pharokka_results/OP310971/OP310971_cds_final_merged_output.tsv | 100 | 20 | 5 |
| LN610577 | ref_le100kb | /Volumes/Extreme_SSD/CRISPR_spacer_analyses/Paper3/pharokka_results/LN610577/LN610577_cds_final_merged_output.tsv | 87 | 3 | 29 |
| PX884080.1 | ref_le100kb | /Volumes/Extreme_SSD/CRISPR_spacer_analyses/Paper3/pharokka_results/PX884080.1/PX884080.1_cds_final_merged_output.tsv | 100 | 16 | 6.25 |
| MN131142 | ref_le100kb | /Volumes/Extreme_SSD/CRISPR_spacer_analyses/Paper3/pharokka_results/MN131142/MN131142_cds_final_merged_output.tsv | 100 | 16 | 6.25 |
| PQ397350 | ref_le100kb | /Volumes/Extreme_SSD/CRISPR_spacer_analyses/Paper3/pharokka_results/PQ397350/PQ397350_cds_final_merged_output.tsv | 49 | 8 | 6.125 |
| JQ067087 | ref_le100kb | /Volumes/Extreme_SSD/CRISPR_spacer_analyses/Paper3/pharokka_results/JQ067087/JQ067087_cds_final_merged_output.tsv | 94 | 3 | 31.3333333 |
| PQ287644 | ref_le100kb | /Volumes/Extreme_SSD/CRISPR_spacer_analyses/Paper3/pharokka_results/PQ287644/PQ287644_cds_final_merged_output.tsv | 243 | 43 | 5.65116279 |
| MK599315 | ref_le100kb | /Volumes/Extreme_SSD/CRISPR_spacer_analyses/Paper3/pharokka_results/MK599315/MK599315_cds_final_merged_output.tsv | 448 | 63 | 7.11111111 |
| PQ738200 | ref_le100kb | /Volumes/Extreme_SSD/CRISPR_spacer_analyses/Paper3/pharokka_results/PQ738200/PQ738200_cds_final_merged_output.tsv | 116 | 13 | 8.92307692 |
| MN304815.1 | ref_le100kb | /Volumes/Extreme_SSD/CRISPR_spacer_analyses/Paper3/pharokka_results/MN304815.1/MN304815.1_cds_final_merged_output.tsv | 53 | 5 | 10.6 |
| ON524861 | ref_le100kb | /Volumes/Extreme_SSD/CRISPR_spacer_analyses/Paper3/pharokka_results/ON524861/ON524861_cds_final_merged_output.tsv | 174 | 9 | 19.3333333 |
| LN889995 | ref_le100kb | /Volumes/Extreme_SSD/CRISPR_spacer_analyses/Paper3/pharokka_results/LN889995/LN889995_cds_final_merged_output.tsv | 53 | 3 | 17.6666667 |
| MN692672 | ref_le100kb | /Volumes/Extreme_SSD/CRISPR_spacer_analyses/Paper3/pharokka_results/MN692672/MN692672_cds_final_merged_output.tsv | 63 | 1 | 63 |
| OR683508 | ref_le100kb | /Volumes/Extreme_SSD/CRISPR_spacer_analyses/Paper3/pharokka_results/OR683508/OR683508_cds_final_merged_output.tsv | 76 | 3 | 25.3333333 |
| BK061475 | ref_le100kb | /Volumes/Extreme_SSD/CRISPR_spacer_analyses/Paper3/pharokka_results/BK061475/BK061475_cds_final_merged_output.tsv | 53 | 6 | 8.83333333 |
| MK813940.1 | ref_le100kb | /Volumes/Extreme_SSD/CRISPR_spacer_analyses/Paper3/pharokka_results/MK813940.1/MK813940.1_cds_final_merged_output.tsv | 105 | 11 | 9.54545455 |
| MG250485 | ref_le100kb | /Volumes/Extreme_SSD/CRISPR_spacer_analyses/Paper3/pharokka_results/MG250485/MG250485_cds_final_merged_output.tsv | 54 | 3 | 18 |
| KU199710 | ref_le100kb | /Volumes/Extreme_SSD/CRISPR_spacer_analyses/Paper3/pharokka_results/KU199710/KU199710_cds_final_merged_output.tsv | 80 | 15 | 5.33333333 |
| PP179321 | ref_le100kb | /Volumes/Extreme_SSD/CRISPR_spacer_analyses/Paper3/pharokka_results/PP179321/PP179321_cds_final_merged_output.tsv | 67 | 1 | 67 |
| MG589385 | ref_le100kb | /Volumes/Extreme_SSD/CRISPR_spacer_analyses/Paper3/pharokka_results/MG589385/MG589385_cds_final_merged_output.tsv | 82 | 9 | 9.11111111 |
| AY962392.1 | ref_le100kb | /Volumes/Extreme_SSD/CRISPR_spacer_analyses/Paper3/pharokka_results/AY962392.1/AY962392.1_cds_final_merged_output.tsv | 278 | 27 | 10.2962963 |
| MK340761 | ref_le100kb | /Volumes/Extreme_SSD/CRISPR_spacer_analyses/Paper3/pharokka_results/MK340761/MK340761_cds_final_merged_output.tsv | 204 | 16 | 12.75 |
| MZ927745 | ref_le100kb | /Volumes/Extreme_SSD/CRISPR_spacer_analyses/Paper3/pharokka_results/MZ927745/MZ927745_cds_final_merged_output.tsv | 59 | 7 | 8.42857143 |
| PQ144888 | ref_le100kb | /Volumes/Extreme_SSD/CRISPR_spacer_analyses/Paper3/pharokka_results/PQ144888/PQ144888_cds_final_merged_output.tsv | 12 | 5 | 2.4 |
| PP496460 | ref_le100kb | /Volumes/Extreme_SSD/CRISPR_spacer_analyses/Paper3/pharokka_results/PP496460/PP496460_cds_final_merged_output.tsv | 55 | 3 | 18.3333333 |
| MK511064 | ref_le100kb | /Volumes/Extreme_SSD/CRISPR_spacer_analyses/Paper3/pharokka_results/MK511064/MK511064_cds_final_merged_output.tsv | 50 | 3 | 16.6666667 |
| MT104473 | ref_le100kb | /Volumes/Extreme_SSD/CRISPR_spacer_analyses/Paper3/pharokka_results/MT104473/MT104473_cds_final_merged_output.tsv | 90 | 3 | 30 |
| OQ925957 | ref_le100kb | /Volumes/Extreme_SSD/CRISPR_spacer_analyses/Paper3/pharokka_results/OQ925957/OQ925957_cds_final_merged_output.tsv | 105 | 15 | 7 |
| PQ287642 | ref_le100kb | /Volumes/Extreme_SSD/CRISPR_spacer_analyses/Paper3/pharokka_results/PQ287642/PQ287642_cds_final_merged_output.tsv | 420 | 55 | 7.63636364 |
| AM910650 | ref_le100kb | /Volumes/Extreme_SSD/CRISPR_spacer_analyses/Paper3/pharokka_results/AM910650/AM910650_cds_final_merged_output.tsv | 79 | 7 | 11.2857143 |
| KM411960 | ref_le100kb | /Volumes/Extreme_SSD/CRISPR_spacer_analyses/Paper3/pharokka_results/KM411960/KM411960_cds_final_merged_output.tsv | 105 | 13 | 8.07692308 |
| PP977452 | ref_le100kb | /Volumes/Extreme_SSD/CRISPR_spacer_analyses/Paper3/pharokka_results/PP977452/PP977452_cds_final_merged_output.tsv | 98 | 16 | 6.125 |
| MK511027 | ref_le100kb | /Volumes/Extreme_SSD/CRISPR_spacer_analyses/Paper3/pharokka_results/MK511027/MK511027_cds_final_merged_output.tsv | 63 | 3 | 21 |
| OR621023 | ref_le100kb | /Volumes/Extreme_SSD/CRISPR_spacer_analyses/Paper3/pharokka_results/OR621023/OR621023_cds_final_merged_output.tsv | 16 | 1 | 16 |
| PP740689 | ref_le100kb | /Volumes/Extreme_SSD/CRISPR_spacer_analyses/Paper3/pharokka_results/PP740689/PP740689_cds_final_merged_output.tsv | 79 | 13 | 6.07692308 |
| OQ067610 | ref_le100kb | /Volumes/Extreme_SSD/CRISPR_spacer_analyses/Paper3/pharokka_results/OQ067610/OQ067610_cds_final_merged_output.tsv | 82 | 5 | 16.4 |
| PQ569625.1 | ref_le100kb | /Volumes/Extreme_SSD/CRISPR_spacer_analyses/Paper3/pharokka_results/PQ569625.1/PQ569625.1_cds_final_merged_output.tsv | 84 | 12 | 7 |
| PV752213.1 | ref_le100kb | /Volumes/Extreme_SSD/CRISPR_spacer_analyses/Paper3/pharokka_results/PV752213.1/PV752213.1_cds_final_merged_output.tsv | 197 | 16 | 12.3125 |
| PV733668.1 | ref_le100kb | /Volumes/Extreme_SSD/CRISPR_spacer_analyses/Paper3/pharokka_results/PV733668.1/PV733668.1_cds_final_merged_output.tsv | 176 | 9 | 19.5555556 |
| OL622073 | ref_le100kb | /Volumes/Extreme_SSD/CRISPR_spacer_analyses/Paper3/pharokka_results/OL622073/OL622073_cds_final_merged_output.tsv | 145 | 5 | 29 |
| MK510986 | ref_le100kb | /Volumes/Extreme_SSD/CRISPR_spacer_analyses/Paper3/pharokka_results/MK510986/MK510986_cds_final_merged_output.tsv | 51 | 9 | 5.66666667 |
| OQ319930 | ref_le100kb | /Volumes/Extreme_SSD/CRISPR_spacer_analyses/Paper3/pharokka_results/OQ319930/OQ319930_cds_final_merged_output.tsv | 62 | 1 | 62 |
| PQ788408 | ref_le100kb | /Volumes/Extreme_SSD/CRISPR_spacer_analyses/Paper3/pharokka_results/PQ788408/PQ788408_cds_final_merged_output.tsv | 77 | 7 | 11 |
| PP003921 | ref_le100kb | /Volumes/Extreme_SSD/CRISPR_spacer_analyses/Paper3/pharokka_results/PP003921/PP003921_cds_final_merged_output.tsv | 87 | 9 | 9.66666667 |
| PP848306 | ref_le100kb | /Volumes/Extreme_SSD/CRISPR_spacer_analyses/Paper3/pharokka_results/PP848306/PP848306_cds_final_merged_output.tsv | 104 | 16 | 6.5 |
| OR683403 | ref_le100kb | /Volumes/Extreme_SSD/CRISPR_spacer_analyses/Paper3/pharokka_results/OR683403/OR683403_cds_final_merged_output.tsv | 55 | 4 | 13.75 |
| HE584812 | ref_le100kb | /Volumes/Extreme_SSD/CRISPR_spacer_analyses/Paper3/pharokka_results/HE584812/HE584812_cds_final_merged_output.tsv | 57 | 3 | 19 |
| OQ658369 | ref_le100kb | /Volumes/Extreme_SSD/CRISPR_spacer_analyses/Paper3/pharokka_results/OQ658369/OQ658369_cds_final_merged_output.tsv | 56 | 5 | 11.2 |
| OR805293 | ref_le100kb | /Volumes/Extreme_SSD/CRISPR_spacer_analyses/Paper3/pharokka_results/OR805293/OR805293_cds_final_merged_output.tsv | 535 | 79 | 6.7721519 |
| KU310943 | ref_le100kb | /Volumes/Extreme_SSD/CRISPR_spacer_analyses/Paper3/pharokka_results/KU310943/KU310943_cds_final_merged_output.tsv | 79 | 11 | 7.18181818 |
| MW329986 | ref_le100kb | /Volumes/Extreme_SSD/CRISPR_spacer_analyses/Paper3/pharokka_results/MW329986/MW329986_cds_final_merged_output.tsv | 81 | 3 | 27 |
| MN871485 | ref_le100kb | /Volumes/Extreme_SSD/CRISPR_spacer_analyses/Paper3/pharokka_results/MN871485/MN871485_cds_final_merged_output.tsv | 107 | 16 | 6.6875 |
| PP541513 | ref_le100kb | /Volumes/Extreme_SSD/CRISPR_spacer_analyses/Paper3/pharokka_results/PP541513/PP541513_cds_final_merged_output.tsv | 202 | 14 | 14.4285714 |
| PV037724 | ref_le100kb | /Volumes/Extreme_SSD/CRISPR_spacer_analyses/Paper3/pharokka_results/PV037724/PV037724_cds_final_merged_output.tsv | 50 | 6 | 8.33333333 |
| MN461279.1 | ref_le100kb | /Volumes/Extreme_SSD/CRISPR_spacer_analyses/Paper3/pharokka_results/MN461279.1/MN461279.1_cds_final_merged_output.tsv | 82 | 5 | 16.4 |
| KY000083 | ref_le100kb | /Volumes/Extreme_SSD/CRISPR_spacer_analyses/Paper3/pharokka_results/KY000083/KY000083_cds_final_merged_output.tsv | 192 | 10 | 19.2 |
| MN871479 | ref_le100kb | /Volumes/Extreme_SSD/CRISPR_spacer_analyses/Paper3/pharokka_results/MN871479/MN871479_cds_final_merged_output.tsv | 111 | 16 | 6.9375 |
| OR805297 | ref_le100kb | /Volumes/Extreme_SSD/CRISPR_spacer_analyses/Paper3/pharokka_results/OR805297/OR805297_cds_final_merged_output.tsv | 89 | 3 | 29.6666667 |
| LN610580 | ref_le100kb | /Volumes/Extreme_SSD/CRISPR_spacer_analyses/Paper3/pharokka_results/LN610580/LN610580_cds_final_merged_output.tsv | 63 | 1 | 63 |
| MK511015 | ref_le100kb | /Volumes/Extreme_SSD/CRISPR_spacer_analyses/Paper3/pharokka_results/MK511015/MK511015_cds_final_merged_output.tsv | 46 | 7 | 6.57142857 |
| OM234793 | ref_le100kb | /Volumes/Extreme_SSD/CRISPR_spacer_analyses/Paper3/pharokka_results/OM234793/OM234793_cds_final_merged_output.tsv | 61 | 11 | 5.54545455 |
| OM131411 | ref_le100kb | /Volumes/Extreme_SSD/CRISPR_spacer_analyses/Paper3/pharokka_results/OM131411/OM131411_cds_final_merged_output.tsv | 56 | 7 | 8 |
| PV844379.1 | ref_le100kb | /Volumes/Extreme_SSD/CRISPR_spacer_analyses/Paper3/pharokka_results/PV844379.1/PV844379.1_cds_final_merged_output.tsv | 112 | 7 | 16 |
| PQ628237 | ref_le100kb | /Volumes/Extreme_SSD/CRISPR_spacer_analyses/Paper3/pharokka_results/PQ628237/PQ628237_cds_final_merged_output.tsv | 86 | 3 | 28.6666667 |
| OM418631 | ref_le100kb | /Volumes/Extreme_SSD/CRISPR_spacer_analyses/Paper3/pharokka_results/OM418631/OM418631_cds_final_merged_output.tsv | 68 | 7 | 9.71428571 |
| MN871484 | ref_le100kb | /Volumes/Extreme_SSD/CRISPR_spacer_analyses/Paper3/pharokka_results/MN871484/MN871484_cds_final_merged_output.tsv | 80 | 7 | 11.4285714 |
| MN602045 | ref_le100kb | /Volumes/Extreme_SSD/CRISPR_spacer_analyses/Paper3/pharokka_results/MN602045/MN602045_cds_final_merged_output.tsv | 63 | 1 | 63 |
| OQ210008 | ref_le100kb | /Volumes/Extreme_SSD/CRISPR_spacer_analyses/Paper3/pharokka_results/OQ210008/OQ210008_cds_final_merged_output.tsv | 196 | 14 | 14 |
| PQ562891 | ref_le100kb | /Volumes/Extreme_SSD/CRISPR_spacer_analyses/Paper3/pharokka_results/PQ562891/PQ562891_cds_final_merged_output.tsv | 65 | 1 | 65 |
| OQ572406 | ref_le100kb | /Volumes/Extreme_SSD/CRISPR_spacer_analyses/Paper3/pharokka_results/OQ572406/OQ572406_cds_final_merged_output.tsv | 35 | 7 | 5 |
| OR621022 | ref_le100kb | /Volumes/Extreme_SSD/CRISPR_spacer_analyses/Paper3/pharokka_results/OR621022/OR621022_cds_final_merged_output.tsv | 20 | 2 | 10 |
| LC765218 | ref_le100kb | /Volumes/Extreme_SSD/CRISPR_spacer_analyses/Paper3/pharokka_results/LC765218/LC765218_cds_final_merged_output.tsv | 430 | 51 | 8.43137255 |
| OQ992204 | ref_le100kb | /Volumes/Extreme_SSD/CRISPR_spacer_analyses/Paper3/pharokka_results/OQ992204/OQ992204_cds_final_merged_output.tsv | 99 | 11 | 9 |
| PV976809.1 | ref_le100kb | /Volumes/Extreme_SSD/CRISPR_spacer_analyses/Paper3/pharokka_results/PV976809.1/PV976809.1_cds_final_merged_output.tsv | 415 | 59 | 7.03389831 |
| PP496470 | ref_le100kb | /Volumes/Extreme_SSD/CRISPR_spacer_analyses/Paper3/pharokka_results/PP496470/PP496470_cds_final_merged_output.tsv | 56 | 3 | 18.6666667 |
| MT108726 | ref_le100kb | /Volumes/Extreme_SSD/CRISPR_spacer_analyses/Paper3/pharokka_results/MT108726/MT108726_cds_final_merged_output.tsv | 102 | 20 | 5.1 |
| PZ122515.1 | ref_le100kb | /Volumes/Extreme_SSD/CRISPR_spacer_analyses/Paper3/pharokka_results/PZ122515.1/PZ122515.1_cds_final_merged_output.tsv | 62 | 3 | 20.6666667 |
| MZ927746 | ref_le100kb | /Volumes/Extreme_SSD/CRISPR_spacer_analyses/Paper3/pharokka_results/MZ927746/MZ927746_cds_final_merged_output.tsv | 56 | 7 | 8 |
| OP556577 | ref_le100kb | /Volumes/Extreme_SSD/CRISPR_spacer_analyses/Paper3/pharokka_results/OP556577/OP556577_cds_final_merged_output.tsv | 64 | 1 | 64 |
| KX458241 | ref_le100kb | /Volumes/Extreme_SSD/CRISPR_spacer_analyses/Paper3/pharokka_results/KX458241/KX458241_cds_final_merged_output.tsv | 51 | 4 | 12.75 |
| OR805295 | ref_le100kb | /Volumes/Extreme_SSD/CRISPR_spacer_analyses/Paper3/pharokka_results/OR805295/OR805295_cds_final_merged_output.tsv | 539 | 83 | 6.4939759 |
| PQ492277 | ref_le100kb | /Volumes/Extreme_SSD/CRISPR_spacer_analyses/Paper3/pharokka_results/PQ492277/PQ492277_cds_final_merged_output.tsv | 58 | 5 | 11.6 |
| KX898400 | ref_le100kb | /Volumes/Extreme_SSD/CRISPR_spacer_analyses/Paper3/pharokka_results/KX898400/KX898400_cds_final_merged_output.tsv | 78 | 5 | 15.6 |
| OQ953987 | ref_le100kb | /Volumes/Extreme_SSD/CRISPR_spacer_analyses/Paper3/pharokka_results/OQ953987/OQ953987_cds_final_merged_output.tsv | 52 | 3 | 17.3333333 |
| MK511029 | ref_le100kb | /Volumes/Extreme_SSD/CRISPR_spacer_analyses/Paper3/pharokka_results/MK511029/MK511029_cds_final_merged_output.tsv | 65 | 3 | 21.6666667 |
| MK511040 | ref_le100kb | /Volumes/Extreme_SSD/CRISPR_spacer_analyses/Paper3/pharokka_results/MK511040/MK511040_cds_final_merged_output.tsv | 75 | 9 | 8.33333333 |
| PP496475 | ref_le100kb | /Volumes/Extreme_SSD/CRISPR_spacer_analyses/Paper3/pharokka_results/PP496475/PP496475_cds_final_merged_output.tsv | 60 | 3 | 20 |
| MK813938.1 | ref_le100kb | /Volumes/Extreme_SSD/CRISPR_spacer_analyses/Paper3/pharokka_results/MK813938.1/MK813938.1_cds_final_merged_output.tsv | 235 | 24 | 9.79166667 |
| MT121968 | ref_le100kb | /Volumes/Extreme_SSD/CRISPR_spacer_analyses/Paper3/pharokka_results/MT121968/MT121968_cds_final_merged_output.tsv | 22 | 3 | 7.33333333 |
| OQ200059 | ref_le100kb | /Volumes/Extreme_SSD/CRISPR_spacer_analyses/Paper3/pharokka_results/OQ200059/OQ200059_cds_final_merged_output.tsv | 196 | 12 | 16.3333333 |
| PP216083 | ref_le100kb | /Volumes/Extreme_SSD/CRISPR_spacer_analyses/Paper3/pharokka_results/PP216083/PP216083_cds_final_merged_output.tsv | 96 | 16 | 6 |
| MW406976 | ref_le100kb | /Volumes/Extreme_SSD/CRISPR_spacer_analyses/Paper3/pharokka_results/MW406976/MW406976_cds_final_merged_output.tsv | 65 | 1 | 65 |
| PP813865 | ref_le100kb | /Volumes/Extreme_SSD/CRISPR_spacer_analyses/Paper3/pharokka_results/PP813865/PP813865_cds_final_merged_output.tsv | 60 | 3 | 20 |
| PP869205 | ref_le100kb | /Volumes/Extreme_SSD/CRISPR_spacer_analyses/Paper3/pharokka_results/PP869205/PP869205_cds_final_merged_output.tsv | 151 | 1 | 151 |
| OQ884027 | ref_le100kb | /Volumes/Extreme_SSD/CRISPR_spacer_analyses/Paper3/pharokka_results/OQ884027/OQ884027_cds_final_merged_output.tsv | 98 | 14 | 7 |
| OQ572408 | ref_le100kb | /Volumes/Extreme_SSD/CRISPR_spacer_analyses/Paper3/pharokka_results/OQ572408/OQ572408_cds_final_merged_output.tsv | 46 | 11 | 4.18181818 |
| AP019535 | ref_le100kb | /Volumes/Extreme_SSD/CRISPR_spacer_analyses/Paper3/pharokka_results/AP019535/AP019535_cds_final_merged_output.tsv | 101 | 14 | 7.21428571 |
| PV565041 | ref_le100kb | /Volumes/Extreme_SSD/CRISPR_spacer_analyses/Paper3/pharokka_results/PV565041/PV565041_cds_final_merged_output.tsv | 148 | 13 | 11.3846154 |
| OL473897 | ref_le100kb | /Volumes/Extreme_SSD/CRISPR_spacer_analyses/Paper3/pharokka_results/OL473897/OL473897_cds_final_merged_output.tsv | 139 | 7 | 19.8571429 |
| KT887558 | ref_le100kb | /Volumes/Extreme_SSD/CRISPR_spacer_analyses/Paper3/pharokka_results/KT887558/KT887558_cds_final_merged_output.tsv | 77 | 5 | 15.4 |
| PP496473 | ref_le100kb | /Volumes/Extreme_SSD/CRISPR_spacer_analyses/Paper3/pharokka_results/PP496473/PP496473_cds_final_merged_output.tsv | 56 | 1 | 56 |
| MN563785 | ref_le100kb | /Volumes/Extreme_SSD/CRISPR_spacer_analyses/Paper3/pharokka_results/MN563785/MN563785_cds_final_merged_output.tsv | 199 | 10 | 19.9 |
| MF975720 | ref_le100kb | /Volumes/Extreme_SSD/CRISPR_spacer_analyses/Paper3/pharokka_results/MF975720/MF975720_cds_final_merged_output.tsv | 62 | 11 | 5.63636364 |
| MK510971 | ref_le100kb | /Volumes/Extreme_SSD/CRISPR_spacer_analyses/Paper3/pharokka_results/MK510971/MK510971_cds_final_merged_output.tsv | 72 | 5 | 14.4 |
| HE983844 | ref_le100kb | /Volumes/Extreme_SSD/CRISPR_spacer_analyses/Paper3/pharokka_results/HE983844/HE983844_cds_final_merged_output.tsv | 81 | 5 | 16.2 |
| PQ533665 | ref_le100kb | /Volumes/Extreme_SSD/CRISPR_spacer_analyses/Paper3/pharokka_results/PQ533665/PQ533665_cds_final_merged_output.tsv | 60 | 3 | 20 |
| LN610578 | ref_le100kb | /Volumes/Extreme_SSD/CRISPR_spacer_analyses/Paper3/pharokka_results/LN610578/LN610578_cds_final_merged_output.tsv | 82 | 5 | 16.4 |
| MF063068 | ref_le100kb | /Volumes/Extreme_SSD/CRISPR_spacer_analyses/Paper3/pharokka_results/MF063068/MF063068_cds_final_merged_output.tsv | 366 | 58 | 6.31034483 |
| MZ955867 | ref_le100kb | /Volumes/Extreme_SSD/CRISPR_spacer_analyses/Paper3/pharokka_results/MZ955867/MZ955867_cds_final_merged_output.tsv | 101 | 11 | 9.18181818 |
| OQ968807 | ref_le100kb | /Volumes/Extreme_SSD/CRISPR_spacer_analyses/Paper3/pharokka_results/OQ968807/OQ968807_cds_final_merged_output.tsv | 100 | 18 | 5.55555556 |
| MF490241 | ref_le100kb | /Volumes/Extreme_SSD/CRISPR_spacer_analyses/Paper3/pharokka_results/MF490241/MF490241_cds_final_merged_output.tsv | 105 | 20 | 5.25 |
| PP704628 | ref_le100kb | /Volumes/Extreme_SSD/CRISPR_spacer_analyses/Paper3/pharokka_results/PP704628/PP704628_cds_final_merged_output.tsv | 57 | 1 | 57 |
| PP496421 | ref_le100kb | /Volumes/Extreme_SSD/CRISPR_spacer_analyses/Paper3/pharokka_results/PP496421/PP496421_cds_final_merged_output.tsv | 51 | 1 | 51 |
| PQ474312 | ref_le100kb | /Volumes/Extreme_SSD/CRISPR_spacer_analyses/Paper3/pharokka_results/PQ474312/PQ474312_cds_final_merged_output.tsv | 107 | 14 | 7.64285714 |
| MZ089728 | ref_le100kb | /Volumes/Extreme_SSD/CRISPR_spacer_analyses/Paper3/pharokka_results/MZ089728/MZ089728_cds_final_merged_output.tsv | 79 | 9 | 8.77777778 |
| OK094665 | ref_le100kb | /Volumes/Extreme_SSD/CRISPR_spacer_analyses/Paper3/pharokka_results/OK094665/OK094665_cds_final_merged_output.tsv | 61 | 11 | 5.54545455 |
| KU199709 | ref_le100kb | /Volumes/Extreme_SSD/CRISPR_spacer_analyses/Paper3/pharokka_results/KU199709/KU199709_cds_final_merged_output.tsv | 57 | 3 | 19 |
| MW651859 | ref_le100kb | /Volumes/Extreme_SSD/CRISPR_spacer_analyses/Paper3/pharokka_results/MW651859/MW651859_cds_final_merged_output.tsv | 61 | 5 | 12.2 |
| KT372691 | ref_le100kb | /Volumes/Extreme_SSD/CRISPR_spacer_analyses/Paper3/pharokka_results/KT372691/KT372691_cds_final_merged_output.tsv | 102 | 16 | 6.375 |
| JX863101 | ref_le100kb | /Volumes/Extreme_SSD/CRISPR_spacer_analyses/Paper3/pharokka_results/JX863101/JX863101_cds_final_merged_output.tsv | 86 | 9 | 9.55555556 |
| PX725670.1 | ref_le100kb | /Volumes/Extreme_SSD/CRISPR_spacer_analyses/Paper3/pharokka_results/PX725670.1/PX725670.1_cds_final_merged_output.tsv | 200 | 18 | 11.1111111 |
| PX674086.1 | ref_le100kb | /Volumes/Extreme_SSD/CRISPR_spacer_analyses/Paper3/pharokka_results/PX674086.1/PX674086.1_cds_final_merged_output.tsv | 63 | 1 | 63 |
| MG432151 | ref_le100kb | /Volumes/Extreme_SSD/CRISPR_spacer_analyses/Paper3/pharokka_results/MG432151/MG432151_cds_final_merged_output.tsv | 87 | 9 | 9.66666667 |
| MH643777 | ref_le100kb | /Volumes/Extreme_SSD/CRISPR_spacer_analyses/Paper3/pharokka_results/MH643777/MH643777_cds_final_merged_output.tsv | 59 | 3 | 19.6666667 |
| PV754031 | ref_le100kb | /Volumes/Extreme_SSD/CRISPR_spacer_analyses/Paper3/pharokka_results/PV754031/PV754031_cds_final_merged_output.tsv | 416 | 45 | 9.24444444 |
| PV400810 | ref_le100kb | /Volumes/Extreme_SSD/CRISPR_spacer_analyses/Paper3/pharokka_results/PV400810/PV400810_cds_final_merged_output.tsv | 235 | 45 | 5.22222222 |
| ON857927 | ref_le100kb | /Volumes/Extreme_SSD/CRISPR_spacer_analyses/Paper3/pharokka_results/ON857927/ON857927_cds_final_merged_output.tsv | 78 | 5 | 15.6 |
| MK510988 | ref_le100kb | /Volumes/Extreme_SSD/CRISPR_spacer_analyses/Paper3/pharokka_results/MK510988/MK510988_cds_final_merged_output.tsv | 74 | 9 | 8.22222222 |
| PP931176 | ref_le100kb | /Volumes/Extreme_SSD/CRISPR_spacer_analyses/Paper3/pharokka_results/PP931176/PP931176_cds_final_merged_output.tsv | 108 | 20 | 5.4 |
| JX434033 | ref_le100kb | /Volumes/Extreme_SSD/CRISPR_spacer_analyses/Paper3/pharokka_results/JX434033/JX434033_cds_final_merged_output.tsv | 56 | 3 | 18.6666667 |
| OR000396 | ref_le100kb | /Volumes/Extreme_SSD/CRISPR_spacer_analyses/Paper3/pharokka_results/OR000396/OR000396_cds_final_merged_output.tsv | 83 | 7 | 11.8571429 |
| MF490239 | ref_le100kb | /Volumes/Extreme_SSD/CRISPR_spacer_analyses/Paper3/pharokka_results/MF490239/MF490239_cds_final_merged_output.tsv | 92 | 1 | 92 |
| JN811560 | ref_le100kb | /Volumes/Extreme_SSD/CRISPR_spacer_analyses/Paper3/pharokka_results/JN811560/JN811560_cds_final_merged_output.tsv | 63 | 5 | 12.6 |
| OR687459 | ref_le100kb | /Volumes/Extreme_SSD/CRISPR_spacer_analyses/Paper3/pharokka_results/OR687459/OR687459_cds_final_merged_output.tsv | 61 | 3 | 20.3333333 |
| OR237807 | ref_le100kb | /Volumes/Extreme_SSD/CRISPR_spacer_analyses/Paper3/pharokka_results/OR237807/OR237807_cds_final_merged_output.tsv | 98 | 9 | 10.8888889 |
| PP337210 | ref_le100kb | /Volumes/Extreme_SSD/CRISPR_spacer_analyses/Paper3/pharokka_results/PP337210/PP337210_cds_final_merged_output.tsv | 210 | 20 | 10.5 |
| PP179316 | ref_le100kb | /Volumes/Extreme_SSD/CRISPR_spacer_analyses/Paper3/pharokka_results/PP179316/PP179316_cds_final_merged_output.tsv | 63 | 5 | 12.6 |
| AM265638 | ref_le100kb | /Volumes/Extreme_SSD/CRISPR_spacer_analyses/Paper3/pharokka_results/AM265638/AM265638_cds_final_merged_output.tsv | 68 | 1 | 68 |
| KX171208 | ref_le100kb | /Volumes/Extreme_SSD/CRISPR_spacer_analyses/Paper3/pharokka_results/KX171208/KX171208_cds_final_merged_output.tsv | 103 | 18 | 5.72222222 |
| MH791399 | ref_le100kb | /Volumes/Extreme_SSD/CRISPR_spacer_analyses/Paper3/pharokka_results/MH791399/MH791399_cds_final_merged_output.tsv | 197 | 10 | 19.7 |
| PV037722 | ref_le100kb | /Volumes/Extreme_SSD/CRISPR_spacer_analyses/Paper3/pharokka_results/PV037722/PV037722_cds_final_merged_output.tsv | 58 | 3 | 19.3333333 |
| MH791403 | ref_le100kb | /Volumes/Extreme_SSD/CRISPR_spacer_analyses/Paper3/pharokka_results/MH791403/MH791403_cds_final_merged_output.tsv | 95 | 1 | 95 |
| OM718002 | ref_le100kb | /Volumes/Extreme_SSD/CRISPR_spacer_analyses/Paper3/pharokka_results/OM718002/OM718002_cds_final_merged_output.tsv | 416 | 53 | 7.8490566 |
| MT119374 | ref_le100kb | /Volumes/Extreme_SSD/CRISPR_spacer_analyses/Paper3/pharokka_results/MT119374/MT119374_cds_final_merged_output.tsv | 99 | 18 | 5.5 |
| MH356729 | ref_le100kb | /Volumes/Extreme_SSD/CRISPR_spacer_analyses/Paper3/pharokka_results/MH356729/MH356729_cds_final_merged_output.tsv | 103 | 5 | 20.6 |
| OQ603352 | ref_le100kb | /Volumes/Extreme_SSD/CRISPR_spacer_analyses/Paper3/pharokka_results/OQ603352/OQ603352_cds_final_merged_output.tsv | 52 | 5 | 10.4 |
| OQ849765 | ref_le100kb | /Volumes/Extreme_SSD/CRISPR_spacer_analyses/Paper3/pharokka_results/OQ849765/OQ849765_cds_final_merged_output.tsv | 47 | 6 | 7.83333333 |
| PV464116 | ref_le100kb | /Volumes/Extreme_SSD/CRISPR_spacer_analyses/Paper3/pharokka_results/PV464116/PV464116_cds_final_merged_output.tsv | 196 | 14 | 14 |
| MN871468 | ref_le100kb | /Volumes/Extreme_SSD/CRISPR_spacer_analyses/Paper3/pharokka_results/MN871468/MN871468_cds_final_merged_output.tsv | 62 | 6 | 10.3333333 |
| MT104465 | ref_le100kb | /Volumes/Extreme_SSD/CRISPR_spacer_analyses/Paper3/pharokka_results/MT104465/MT104465_cds_final_merged_output.tsv | 50 | 1 | 50 |
| MT118296 | ref_le100kb | /Volumes/Extreme_SSD/CRISPR_spacer_analyses/Paper3/pharokka_results/MT118296/MT118296_cds_final_merged_output.tsv | 47 | 1 | 47 |
| OR801193 | ref_le100kb | /Volumes/Extreme_SSD/CRISPR_spacer_analyses/Paper3/pharokka_results/OR801193/OR801193_cds_final_merged_output.tsv | 28 | 6 | 4.66666667 |
| MT711889 | ref_le100kb | /Volumes/Extreme_SSD/CRISPR_spacer_analyses/Paper3/pharokka_results/MT711889/MT711889_cds_final_merged_output.tsv | 56 | 3 | 18.6666667 |
| MK511028 | ref_le100kb | /Volumes/Extreme_SSD/CRISPR_spacer_analyses/Paper3/pharokka_results/MK511028/MK511028_cds_final_merged_output.tsv | 63 | 3 | 21 |
| OQ319931 | ref_le100kb | /Volumes/Extreme_SSD/CRISPR_spacer_analyses/Paper3/pharokka_results/OQ319931/OQ319931_cds_final_merged_output.tsv | 56 | 5 | 11.2 |
| MZ089731 | ref_le100kb | /Volumes/Extreme_SSD/CRISPR_spacer_analyses/Paper3/pharokka_results/MZ089731/MZ089731_cds_final_merged_output.tsv | 79 | 9 | 8.77777778 |
| MT118304 | ref_le100kb | /Volumes/Extreme_SSD/CRISPR_spacer_analyses/Paper3/pharokka_results/MT118304/MT118304_cds_final_merged_output.tsv | 58 | 9 | 6.44444444 |
| KT734862 | ref_le100kb | /Volumes/Extreme_SSD/CRISPR_spacer_analyses/Paper3/pharokka_results/KT734862/KT734862_cds_final_merged_output.tsv | 95 | 1 | 95 |
| MH973725 | ref_le100kb | /Volumes/Extreme_SSD/CRISPR_spacer_analyses/Paper3/pharokka_results/MH973725/MH973725_cds_final_merged_output.tsv | 78 | 5 | 15.6 |
| MH791408 | ref_le100kb | /Volumes/Extreme_SSD/CRISPR_spacer_analyses/Paper3/pharokka_results/MH791408/MH791408_cds_final_merged_output.tsv | 199 | 10 | 19.9 |
| OQ210007 | ref_le100kb | /Volumes/Extreme_SSD/CRISPR_spacer_analyses/Paper3/pharokka_results/OQ210007/OQ210007_cds_final_merged_output.tsv | 192 | 12 | 16 |
| KC862295 | ref_le100kb | /Volumes/Extreme_SSD/CRISPR_spacer_analyses/Paper3/pharokka_results/KC862295/KC862295_cds_final_merged_output.tsv | 173 | 11 | 15.7272727 |
| JX403939 | ref_le100kb | /Volumes/Extreme_SSD/CRISPR_spacer_analyses/Paper3/pharokka_results/JX403939/JX403939_cds_final_merged_output.tsv | 77 | 7 | 11 |
| PQ790658 | ref_le100kb | /Volumes/Extreme_SSD/CRISPR_spacer_analyses/Paper3/pharokka_results/PQ790658/PQ790658_cds_final_merged_output.tsv | 76 | 5 | 15.2 |
| PQ741806 | ref_le100kb | /Volumes/Extreme_SSD/CRISPR_spacer_analyses/Paper3/pharokka_results/PQ741806/PQ741806_cds_final_merged_output.tsv | 413 | 51 | 8.09803922 |
| OQ319932 | ref_le100kb | /Volumes/Extreme_SSD/CRISPR_spacer_analyses/Paper3/pharokka_results/OQ319932/OQ319932_cds_final_merged_output.tsv | 102 | 14 | 7.28571429 |
| PV105456 | ref_le100kb | /Volumes/Extreme_SSD/CRISPR_spacer_analyses/Paper3/pharokka_results/PV105456/PV105456_cds_final_merged_output.tsv | 196 | 12 | 16.3333333 |
| KX066068 | ref_le100kb | /Volumes/Extreme_SSD/CRISPR_spacer_analyses/Paper3/pharokka_results/KX066068/KX066068_cds_final_merged_output.tsv | 56 | 5 | 11.2 |
| OR733707 | ref_le100kb | /Volumes/Extreme_SSD/CRISPR_spacer_analyses/Paper3/pharokka_results/OR733707/OR733707_cds_final_merged_output.tsv | 106 | 14 | 7.57142857 |
| MN615698 | ref_le100kb | /Volumes/Extreme_SSD/CRISPR_spacer_analyses/Paper3/pharokka_results/MN615698/MN615698_cds_final_merged_output.tsv | 67 | 1 | 67 |
| PQ758388 | ref_le100kb | /Volumes/Extreme_SSD/CRISPR_spacer_analyses/Paper3/pharokka_results/PQ758388/PQ758388_cds_final_merged_output.tsv | 121 | 9 | 13.4444444 |
| PQ493644.1 | ref_le100kb | /Volumes/Extreme_SSD/CRISPR_spacer_analyses/Paper3/pharokka_results/PQ493644.1/PQ493644.1_cds_final_merged_output.tsv | 63 | 1 | 63 |
| MK511032 | ref_le100kb | /Volumes/Extreme_SSD/CRISPR_spacer_analyses/Paper3/pharokka_results/MK511032/MK511032_cds_final_merged_output.tsv | 71 | 3 | 23.6666667 |

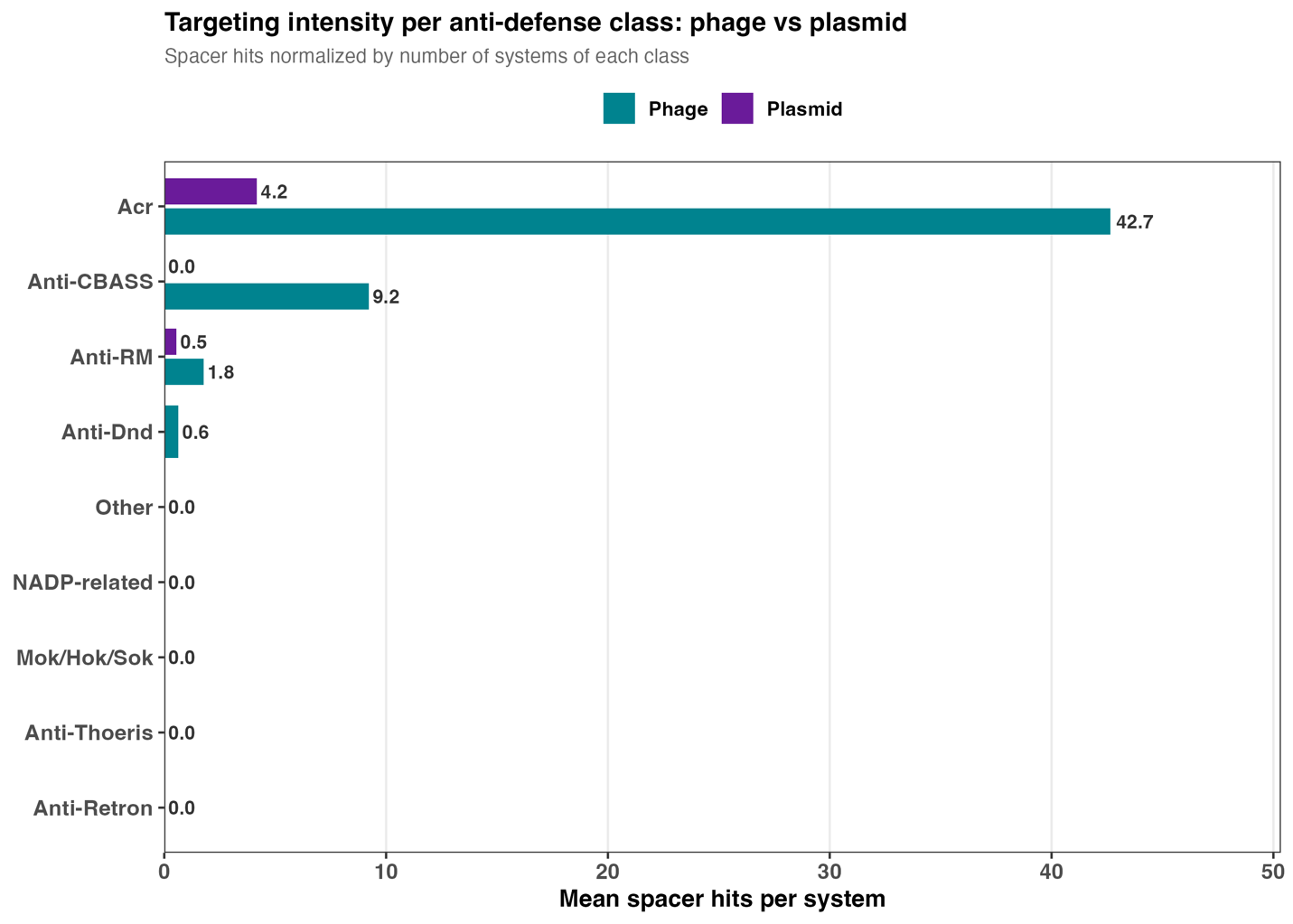

SI Figure 1. Anti-defense targeting intensity by system class, phage versus plasmid. Mean spacer hits per annotated system for each anti-defense class, shown separately for phage (teal) and plasmid (purple) genomes. Acr systems are the most intensively targeted class in both mobile-element types (42.7 hits per system in phages; 4.2 in plasmids), far exceeding all other anti-defense classes. Classes with no targeted systems in a given mobile-element type are shown as 0.0. Values correspond to SI Table 1.

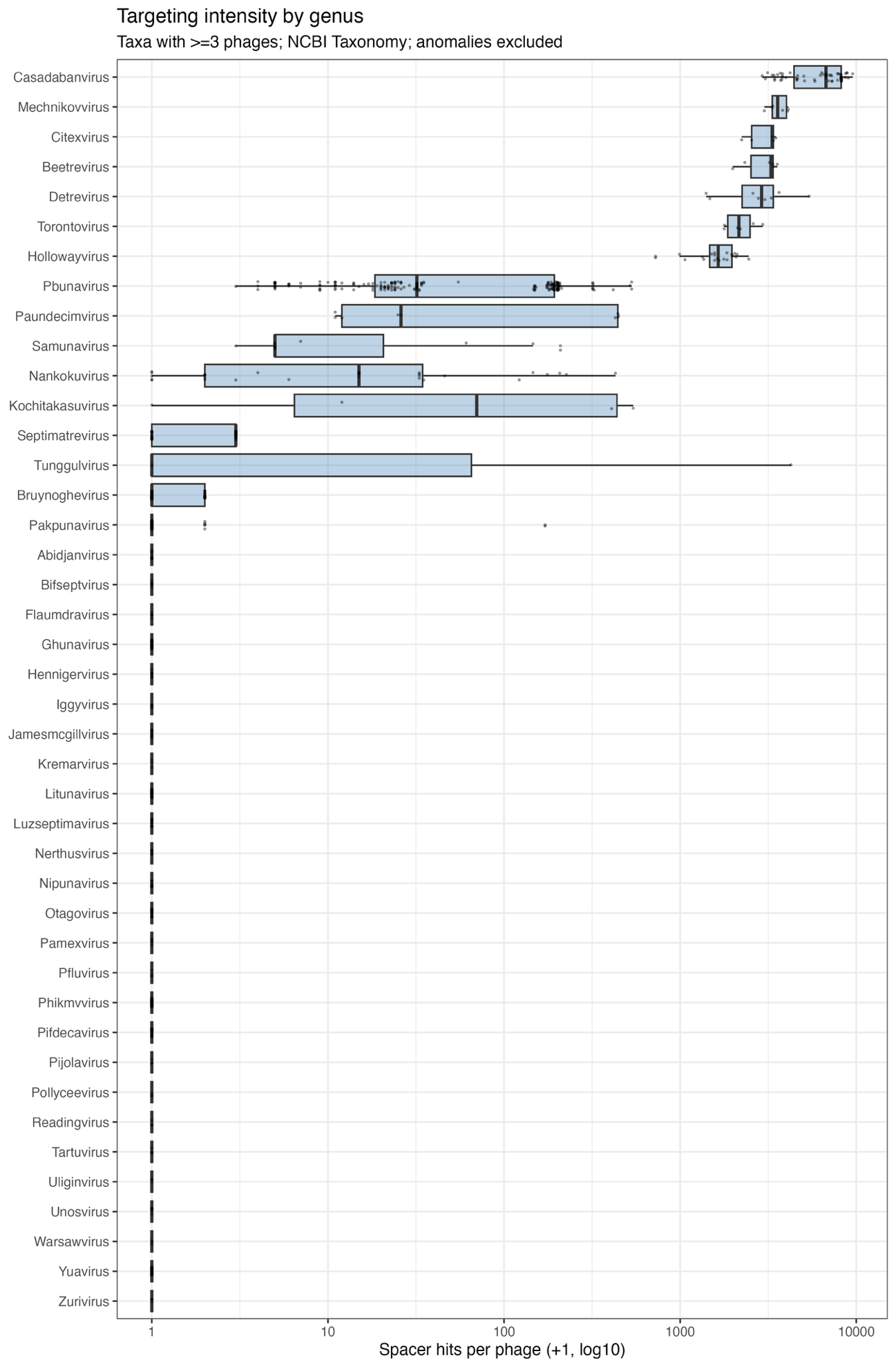

SI Figure 2. Per-genus spacer-targeting intensity across phage genera. Distribution of spacer hits per phage (log₁₀ scale, +1 pseudocount) for each phage genus with ≥3 classified genomes; boxes show the interquartile range and median, whiskers the range. Genera are ordered by targeting intensity. Targeting varies by several orders of magnitude across genera, with Casadabanvirus the most intensively targeted. Underlying per-genus statistics are given in SI Table 2.

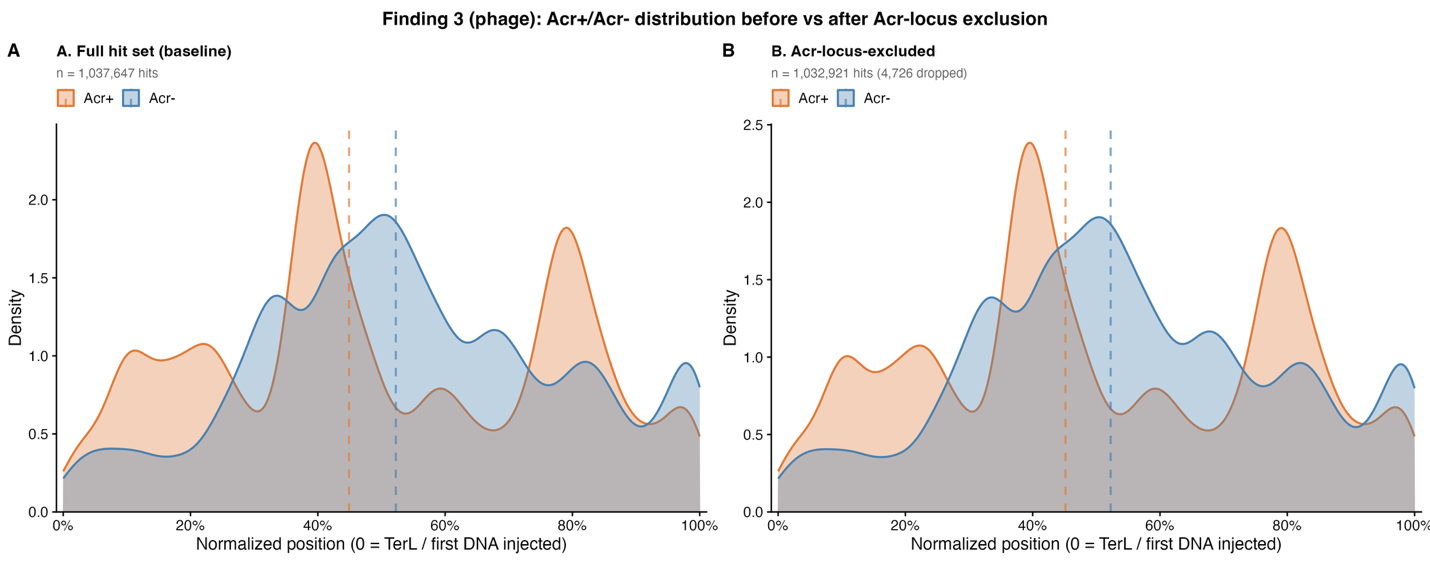

SI Figure 3. The Acr+ spacer-hit positional shift is not driven by spacers directly targeting Acr loci in phages. Kernel density of normalized spacer-hit positions (0 = TerL/first-injected end) for Acr+ (orange) and Acr- (blue) phages, with group medians (dashed lines). (A) All retained spacer hits (n = 1,037,647). (B) After removing spacer hits whose midpoint falls within annotated Anti-CRISPR system coordinates (n = 1,032,921; 4,726 hits removed). The earlier-targeting of Acr+ phages persists after excluding direct Acr-locus hits, indicating a genome-wide positional shift rather than an artifact of direct Acr targeting.

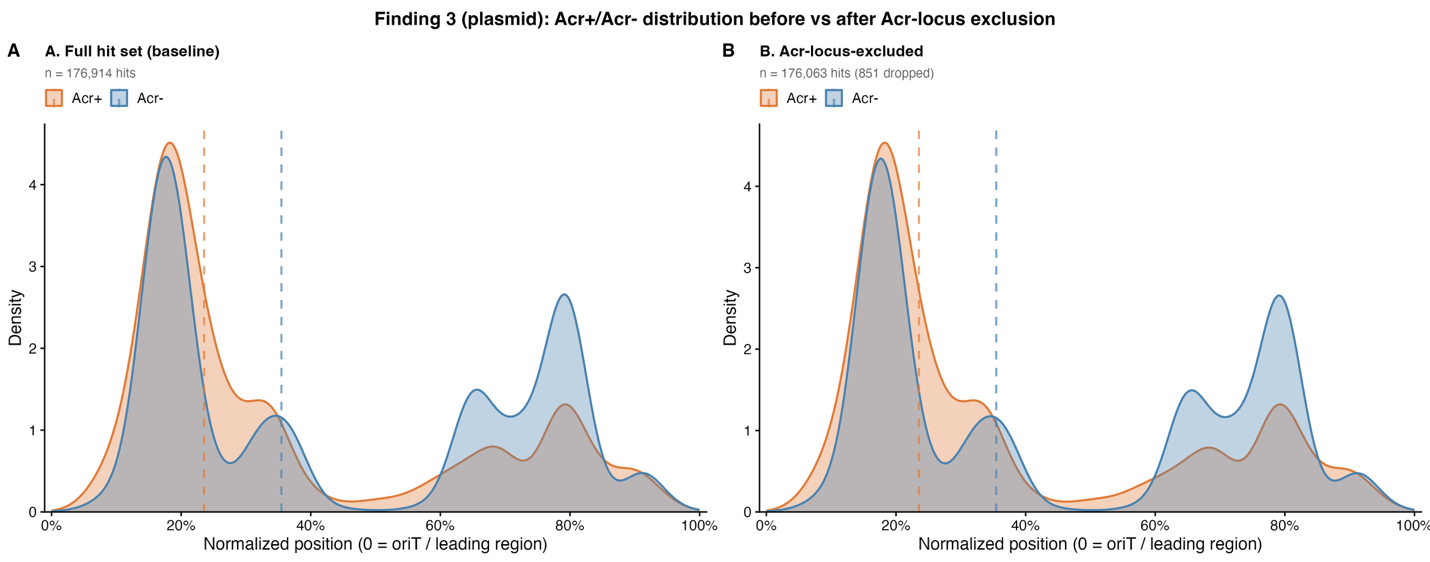

SI Figure 4. The Acr+ spacer-hit positional shift is not driven by spacers directly targeting Acr loci in plasmids. Kernel density of normalized spacer-hit positions (0 = oriT/leading region) for Acr+ (orange) and Acr− (blue) plasmids, with group medians (dashed lines). (A) All retained spacer hits (n = 176,914). (B) After removing spacer hits overlapping annotated Anti-CRISPR system coordinates (n = 8,565; 223 hits removed). As in phages, the positional shift persists after excluding direct Acr-locus hits.
